# Structure-guided design of a cell penetrating peptide preventing cAMP modulation of HCN channels

**DOI:** 10.1101/253096

**Authors:** Andrea Saponaro, Francesca Cantini, Alessandro Porro, Annalisa Bucchi, Dario Di Francesco, Vincenzo Maione, Chiara Donadoni, Bianca Introini, Pietro Mesirca, Matteo E. Mangoni, Gerhard Thiel, Lucia Banci, Bina Santoro, Anna Moroni

## Abstract

The auxiliary subunit TRIP8b prevents cAMP activation of HCN channels by antagonizing its binding to their cyclic-nucleotide binding domain (CNBD). By determining an NMR-derived structure of the complex formed by the HCN2 channel CNBD and a minimal TRIP8b fragment, TRIP_nano_, we show here a bipartite interaction between the peptide and CNBD which prevents cAMP binding in two ways: through direct competition for binding at the distal C-helix of the CNBD; and through an allosteric reduction in cAMP affinity induced by TRIP8b binding to the CNBD N-bundle loop. TRIP_nano_ abolishes cAMP binding in all three isoforms, HCN1, HCN2 and HCN4 and can be used to prevent cAMP stimulation in native f-channels. Application of TRIP8b_nano_, or its delivery via a cell-penetrating sequence, in sinoatrial node myocytes, selectively inhibits beta-adrenergic stimulation of the native I_f_ current and mimics the physiological concentrations of acetylcholine leading to a 30% reduction in the spontaneus rate of action potential firing.

## Introduction

Hyperpolarization-activated cyclic nucleotide-regulated (HCN1-4) channels are the molecular correlate of the I_f_/I_h_ current, which plays a key role in controlling several higher order electrophysiological functions, including dendritic integration and intrinsic rhythmicity both in cardiac and neuronal cells (Robinson & Siegelbaum, 2003). Unique among the voltage-gated ion channel superfamily, HCN channels are modulated by the direct binding of cAMP to their C-terminal CNBD. cAMP binding enhances channel opening upon hyperpolarization via conformational changes in the CNBD that are propagated, through the C-linker domain, to the pore (Wainger *et al*, 2001; Zagotta *et al*, 2003).

In addition to cAMP, HCN channels are regulated by TRIP8b, their brain-specific auxiliary (β) subunit, which modulates channel trafficking and gating (Santoro *et al*, 2009; Zolles *et al*, 2009). TRIP8b binds HCN channels in two distinct sites: the tetratricopeptide repeat (TPR) domain, which binds the last three amino acids (SNL) of HCN channels; and the TRIP8b_core_ domain, which interacts with the HCN channel CNBD domain (Santoro *et al*, 2011). Although TRIP8b is subject to alternative splicing giving rise to nine isoforms with different effects on HCN trafficking (Santoro *et al*, 2009; Piskorowski *et al*, 2011; Lewis *et al*, 2009), all isoforms act in the same way on the electrical activity of HCN channels: they bind to the cAMP-free state of the CNBD and thus antagonize the effect of the ligand on the voltage dependency of the channel (Hu *et al*, 2013).

In previous work, we contributed to elucidating the dual mechanism of action of cAMP and TRIP8b on HCN channels by using a simplified system that included the HCN2 CNBD and the TRIP8b_core_ protein fragments (Saponaro *et al*, 2014). By means of solution NMR spectroscopy we described in atomic detail the cAMP-induced conformational changes occurring in the HCN channel CNBD. These include a major rearrangement of the C-terminal helix of the CNBD (C-helix), which undergoes lateral translation and folding upon cAMP binding; and an upward movement of the N-terminal helical bundle, a helix-turn-helix motif immediately upstream of the CNBD β-roll, which presumably initiates the cAMP-induced facilitation of pore opening. These movements are ostensibly inhibited by TRIP8b binding to both the C-helix and N-terminal helical bundle, thus explaining the allosteric inhibition exerted by TRIP8b on the HCN channel response to cAMP (Hu *et al*, 2013). The above TRIP8b interaction sites were validated by deletion analysis, confirming that both the N-terminal helical bundle and the C-helix are necessary but not sufficient for TRIP8b_core_ binding to the CNBD (Saponaro *et al*, 2014). Despite these findings, several reports have suggested a direct competition mechanism between the two ligands for a common binding region that comprises the C-helix and the phosphate binding cassette, PBC, in the HCN channel CNBD (Han *et al*, 2011; Deberg *et al*, 2015; Bankston *et al*, 2017).

A comprehensive structural model of the complex between TRIP8b and the HCN channel CNBD, which may explain the interaction in atomic detail, is therefore crucial for solving the apparent discrepancies between the two proposed models.

We present here a NMR-based 3D model structure of the complex formed by a minimal TRIP8b fragment and the CNBD of the human HCN2 channel isoform, which provides molecular support for both direct and indirect (allosteric) competition modes exerted by TRIP8b on cAMP binding. TRIP8b_nano_ prevented cAMP stimulation in all HCN isoform tested (HCN1, 2 and 4) and was further employed to manipulate the native I_f_ current in cardiac pacemaker cells. This result opens the possibility of controlling *in vivo* the cAMP-dependent facilitation of HCN channel opening, which represents the basis for the autonomic regulation of cardiac activity and spontaneous neuronal firing (Robinson & Siegelbaum, 2003).

## Results

We have previously shown that TRIP8b_core_ (residues 223 − 303 of mouse TRIP8b splice variant 1a4, hereafter TRIP8b) interacts with two elements of the isolated CNBD protein fragment from HCN channels (residues 521 - 672 of human HCN2, hereafter CNBD): the C-helix and the N-bundle loop, a sequence connecting helix E’ of the C-linker with helix A of the CNBD (Saponaro *et al*, 2014). Biochemical assays confirmed that each of these two elements i.e. the N-bundle loop and C-helix is necessary but not sufficient for binding (Saponaro *et al*, 2014).

To understand the interaction in atomic detail, we used solution NMR spectroscopy to characterize the structural properties of the CNBD-TRIP8b_core_ complex. However, the NMR spectra of TRIP8b_core_ showed very few signals. In order to improve the quality of the NMR spectra, we reduced the length of the TRIP8b fragment by progressively removing residues at the N- and C-termini with no predicted secondary structure. The truncated peptides were then tested for CNBD binding activity by isothermal titration calorimetry (ITC). We thus identified a 40 amino acid peptide (TRIP8b_nano_, comprising residues 235 – 275 of TRIP8b, Fig 1A) with a binding K_D_ of 1.5 ± 0.1 µM, a value very similar to the K_D_ of 1.2 ± 0.1 µM obtained with TRIP8b_core_ (Fig 1B). TRIP8b_nano_ was therefore employed for all subsequent NMR experiments, resulting in a remarkable improvement in the spectral quality and sample stability.

**Figure 1.**
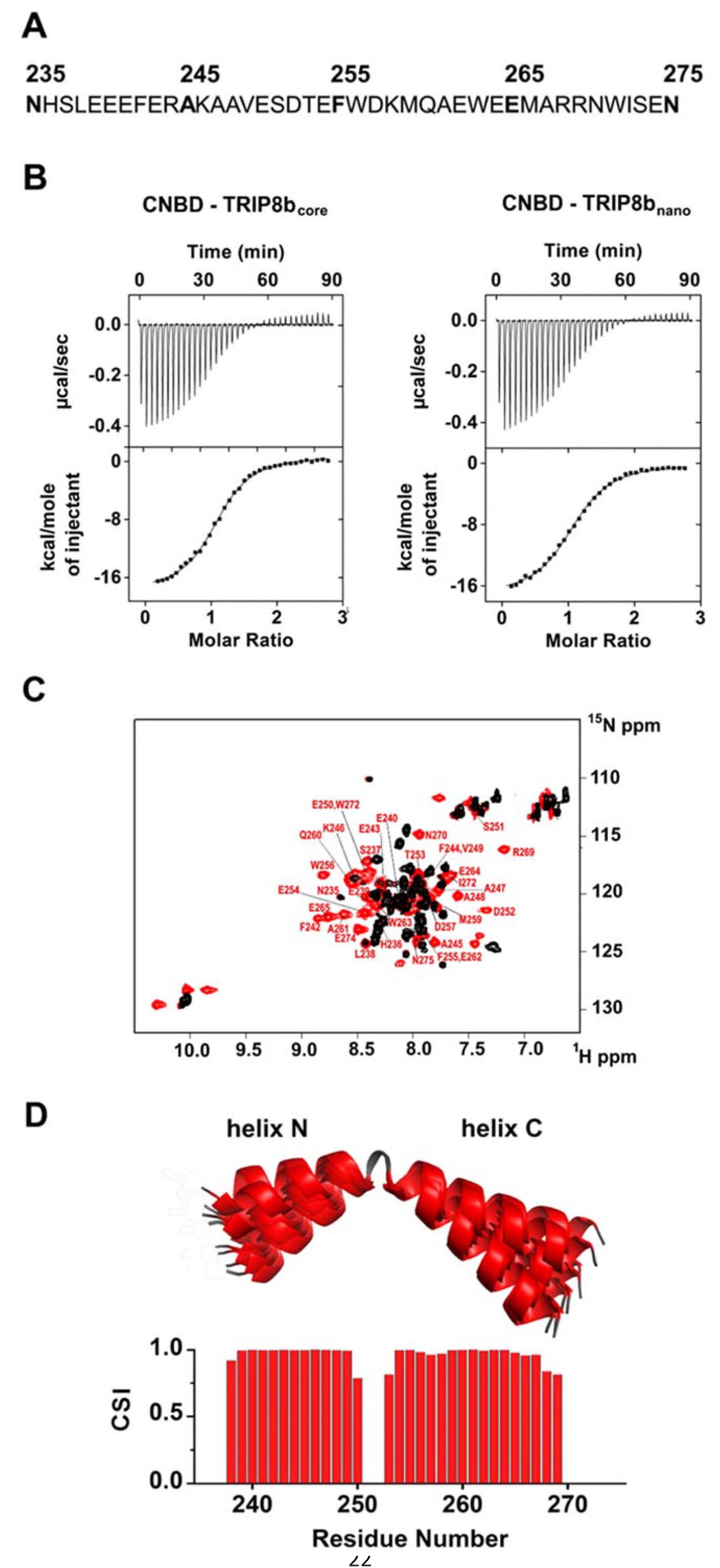
Functional and structural characterization of TRIP8b_nano_. A Primary sequence of TRIP8b_nano_. Amino acid numbering refers to full length mouse TRIP8b (1a4). B Binding of TRIP8b_core_ and TRIP8b_nano_ to purified His_6_-MBP-CNBD measured by Isothermal titration calorimetry (ITC). Upper panel, heat changes (μcal/sec) during successive injections of 8 μL of the corresponding TRIP8b peptide (200 μM) into the chamber containing His_6_-MBP-CNBD (20 μM). Lower panel, binding curve obtained from data displayed in the upper panel. The peaks were integrated, normalized to TRIP8b peptide concentration, and plotted against the molar ratio (TRIP8b peptide / His_6_-MBP-CNBD). Solid line represents a nonlinear least-squares fit to a single-site binding model, yielding, in the present examples, a K_D_ = 1.2 ± 0.1 μM for TRIP8b_core_ and K_D_ = 1.4 ± 0.1 μM for TRIP8b_nano_. C Evidence for TRIP8b_nano_ folding upon CNBD binding based on the superimposition of the [^1^H, ^15^N] heteronuclear single quantum coherence (HSQC) NMR spectrum of CNBD-free TRIP8b_nano_ (black) and CNBD-bound TRIP8b_nano_ (red). The latter experiment was performed at the molar ratio ([CNBD]/[ TRIP8b_nano_]) = 3. The backbone amide (HN) signals of the residues of CNBD-bound TRIP8b_nano_ are labelled in red. D (Top) Ribbon representation of the 10 lowest energy conformers of TRIP8b_nano_ bound to CNBD used for *in silico* modelling of CNBD-TRIP8b_nano_ complex. The unfolded regions at the N- and C-termini of the construct (residues 235–237 and 270 – 275) are omitted for clarity. (Bottom) Chemical Shift Index (CSI, calculated using TALOS+) plotted as a function of the residue number of TRIP8b_nano_ bound to CNBD. Positive values represent helical propensity.

### Structural characterization of TRIP8b_nano_ bound to CNBD

The comparison of the ^1^H-^15^N HSQC spectra of TRIP8b_nano_ with and without CNBD bound shows that the peptide folds upon interaction with the CNBD. Thus, ^1^H-^15^N HSQC spectrum of TRIP8b_nano_ without CNBD shows a limited ^1^H resonance dispersion characteristic of unstructured proteins (Dyson & Wright, 2004), while a larger number of well-dispersed amide signals appear in the spectrum of the CNBD-bound form (Fig 1C). Importantly, we were now able to assign the backbone chemical shift resonances of TRIP8b_nano_ bound to the CNBD. The φ and ψ dihedral angles obtained from the NMR assignment indicate that the peptide displays two α-helices (stretch L_238_-E_250_ named helix N and stretch T_253_-R_269_ named helix C) when bound to CNBD. The helices are separated by two amino acids; three and six residues at the N- and C- termini, respectively, are unstructured (Fig 1D).

### Structural characterization of CNBD bound to TRIP8b_nano_

NMR-analysis of the CNBD fragment bound to TRIP8b_nano_ revealed that the interaction with the peptide does not affect the overall fold of the protein. Thus, the CNBD adopts the typical fold of the cAMP-free state, in line with previous evidence that this is the form bound by TRIP8b (Deberg *et al*, 2015; Saponaro *et al*, 2014). More specifically, the secondary structure elements of the cAMP-free CNBD are all conserved in the TRIP8b_nano_–bound CNBD (Fig 2). This finding generally agrees with a recent double electron-electron resonance (DEER) analysis of the CNBD-TRIP8b interaction, which showed that TRIP8b binds to a conformation largely similar to the cAMP-free state (Deberg *et al*, 2015). Despite the overall agreement with the DEER study, the NMR data also reveal a new and unexpected feature of TRIP8b binding to the CNBD. Our results surprisingly show that TRIP8b_nano_ induces, upon binding to the CNBD, a well-defined secondary structure of the distal region of the C-helix (Fig 2). This means that the distal region of the C-helix (residues 657-662), which is unstructured in the free form of the CNBD (Saponaro *et al*, 2014; Lee & MacKinnon, 2017), extends into a helical structure upon ligand binding irrespectively of whether the ligand is cAMP (Saponaro *et al*, 2014; Puljung & Zagotta, 2013; Lee & MacKinnon, 2017) or TRIP8b (Fig 2). In contrast, and very differently from cAMP, which directly contacts the P-helix in the Phosphate Binding cassette (PBC) and causes its folding (Saponaro *et al*, 2014; Lee & MacKinnon, 2017), the NMR data show that TRIP8b_nano_ binding to the CNBD does not induce P-helix formation (Fig 2).

**Figure 2.**
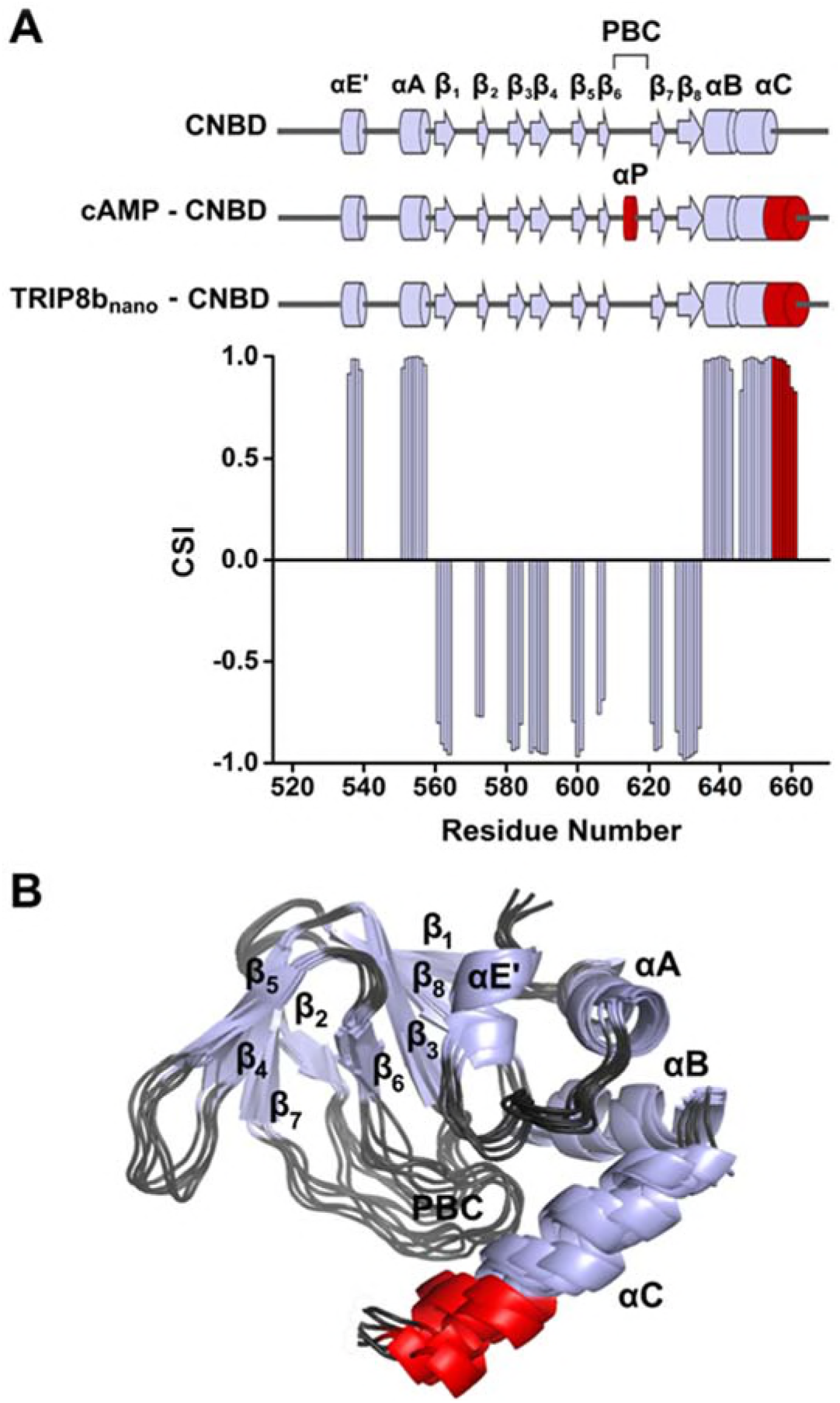
NMR structure of CNBD bound to TRIP8b_nano_. A (Top) comparison of secondary structure elements of cAMP-free CNBD (Saponaro *et al*, 2014), cAMP-bound CNBD (Zagotta *et al*, 2003) and cAMP-free CNBD bound to TRIP8b_nano_ (this study). Secondary structure elements are indicated by arrows (β-strands) and cylinders (α-helices) and labelled. The loop between β_6_ and β_7_ constitutes the Phosphate Binding Cassette (PBC). The elements that fold upon binding of cAMP and TRIP8b_nano_ are shown in red. (Bottom) Chemical Shift Index (CSI, calculated using TALOS+) plotted as a function of the residue number of CNBD bound to TRIP8b_nano_. Positive values represent helical propensity, while negative values represent strands. B Ribbon representation of the 10 lowest energy conformers of CNBD bound to TRIP8b_nano_ used for *in silico* modelling of CNBD-TRIP8b_nano_ complex. Secondary structure elements are coloured in light gray and labelled. Loop regions are coloured in dark gray. The distal region of the C-helix (residues 657-662), which is unfolded in the free form of the CNBD (Saponaro *et al*, 2014) and folds upon TRIP8b_nano_ binding, is coloured in red. The unfolded regions at the N- and C-termini of the construct (residues 521–532 and 663 – 672 respectively) are omitted for clarity.

### Modelling the CNBD-TRIP8b_nano_ complex

Despite the significant improvement in sample stability and NMR spectra quality achieved upon TRIP8b_nano_ binding, we were still unable to assign the side chains of both proteins in the complex and thus could not solve the solution structure of the complex by the canonical NMR procedure. We therefore built a model of the CNBD-TRIP8b_nano_ complex by docking the two NMR-derived structures using the Haddock program (a detailed description of how the respective structures were generated is provided in Materials and Methods and Appendix Table S1).

In order to define the active residues (ambiguous interaction restraints) on the CNBD we used the chemical shift perturbation values as described in Appendix Fig S1. For TRIP8b_nano_, we defined as active a stretch of residues, E_239_-E_243_, previously identified as critical for the interaction (Santoro *et al*, 2009, 2011). Output clusters of this first molecular docking calculation (settings can be found in Material and Methods) were further screened for TRIP8b_nano_ orientations in agreement with a previous DEER analysis, which placed TRIP8b residue A_248_ closer to the proximal portion and TRIP8b residue A_261_ closer to the distal portion of the CNBD C-helix (Deberg *et al*, 2015). Remarkably, in all clusters thus selected, residues E_264_ or E_265_ in TRIP8b were found to interact with residues K_665_ or K_666_ of the CNBD (Appendix Fig S2). This finding was notable, because we previously identified K_665_/K_666_ as being critical for TRIP8b interaction in a biochemical binding assay (Saponaro *et al*, 2014). We thus proceeded to individually mutate each of these four positions, and test their effect on binding affinity through ITC. As expected, reverse charge mutations K_665_E or K_666_E (CNBD) as well as E_264_K or E_265_K (TRIP8b_nano_) each strongly reduced the CNBD/TRIP8b_nano_ binding affinity (Appendix Fig S3).

Based on these observations, we performed a second molecular docking calculation, including E_264_ and E_265_ as additional active residues for TRIP8b_nano_. This procedure resulted in the model shown in Fig 3, which represents the top-ranking cluster for energetic and scoring function (Appendix Table S2) and was fully validated by mutagenesis analysis as described below. Scrutiny of the model shows that TRIP8b_nano_ binds to both the C-helix and the N-bundle loop (Fig 3A). Binding to the C-helix is mainly guided by electrostatic interactions between the negative charges on TRIP8b_nano_, and the positive charges on the CNBD (Fig 3A). As shown in Fig 3B, the model highlights a double saline bridge (K_665_ and K_666_ of CNBD with E_265_ and E_264_ of TRIP8b_nano_) in line with the ITC results described above (Appendix Fig S3). Of note, the contribution of residue R_662_ to the binding is also consistent with previous experiments showing residual TRIP8b interaction in a CNBD deletion mutant ending at position 663 (Saponaro *et al*, 2014). Our modelling data suggest that, upon folding of the distal portion of the C-helix, the side chains of residues R_662_ and R_665_ face to the inside when contacting cAMP but face to the outside when binding TRIP8b (Appendix Fig S4).

In addition to clarifying the role of residues in the distal portion of the CNBD C-helix, the model also highlights a second important cluster of electrostatic interactions with R_650_ in the proximal portion of the CNBD C-helix contacting E_240_ and E_241_ in helix N of TRIP8b_nano_ (Fig 3C). To confirm the contribution of these residues, we reversed charges and tested each residue mutation for binding in ITC. The results in Appendix Fig S3 show that R_650_E caused a more than six-fold reduction in binding affinity for TRIP8b_nano_, with smaller but significant effects seen also for E_240_R and E_241_R.

**Figure 3.**
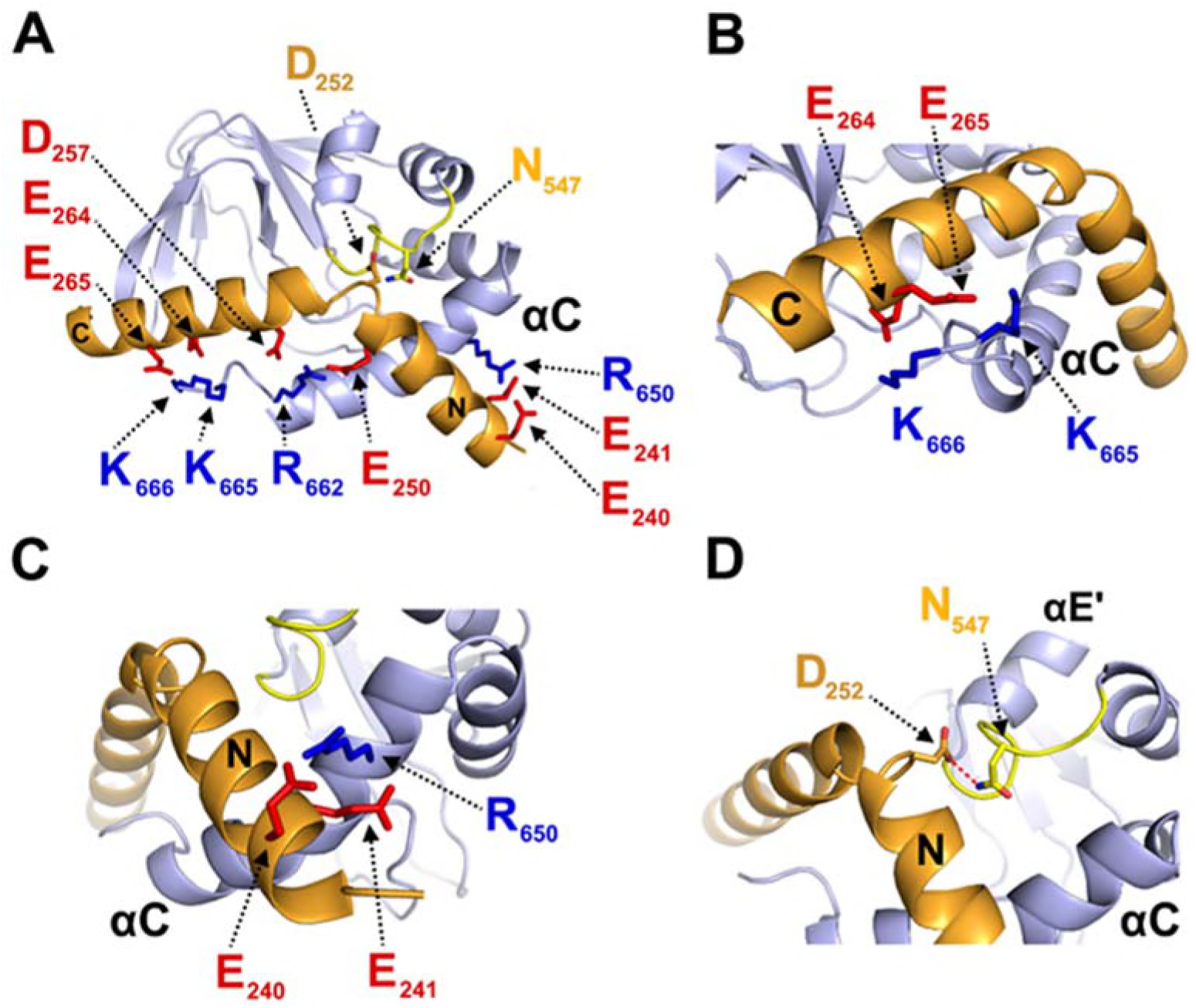
Structural model of CNBD–TRIP8b_nano_ complex. A Ribbon representation of the complex where CNBD is in gray and TRIP8b_nano_ is in orange. Helix N (N) and helix C (C) of TRIP8b_nano_ are labelled. C-helix of CNBD (αC) is labelled, while N-bundle loop is coloured in yellow. Positively charged residues of C-helix CNBD (blue) and negatively charged residues of TRIP8b_nano_ (red) involved in salt bridges are shown as sticks and labelled. N_547_ of the N-bundle loop (yellow) and D_252_ of TRIP8b_nano_ (orange) are shown as sticks and labelled. B Close view of K_665_ and K_666_ of CNBD that interact respectively with E_265_ and E_264_ of TRIP8b_nano_. C-helix (αC) of CNBD, and Helix C (C) of TRIP8b_nano_ are labelled. C Close view of R_650_ of CNBD that is positioned between E_240_ and E_241_ of TRIP8b_nano_. C-helix (αC) of CNBD, and Helix N (N) of TRIP8b_nano_ are labelled. D Close view of N_547_ of N-bundle loop that forms a hydrogen bond (red dashed line) with D_252_ of TRIP8b_nano_. Helix E’ (αE’) and C-helix (αC) of CNBD, and Helix N (N) of TRIP8b_nano_ are labelled.

A third important contact highlighted by the model is the interaction between N_547_ in the N-bundle loop of the CNBD and D_252_ in the link between helix N and helix C of TRIP8b_nano_ (Fig 3D). We tested this potential interaction by disrupting the expected hydrogen bond between N_547_ and the carboxyl group of the negative residue (D_252_) in TRIP8b_nano_. The asparagine in CNBD was mutated into aspartate (N_547_D) to generate an electrostatic repulsion for D_252_, and the carboxyl group in D_252_ of TRIP8b_nano_ was removed by mutation into asparagine (D_252_N). As predicted, N547D greatly reduced binding to TRIP8b in ITC assays (Appendix Fig S3), with a smaller but significant effect observed also for D_252_N (Appendix Fig S3). These results confirm and extend our previous finding that the N-bundle loop contributes in a substantial manner to the binding of TRIP8b (Saponaro *et al*, 2014).

### TRIP8b_nano_ as a tool for the direct regulation of native HCN currents

Next, we asked whether the relatively short TRIP8b_nano_ peptide could be used to block the response of HCN channels to cAMP by delivering the peptide to full length channels heterologusly expressed in cells. To this end, we dialyzed TRIP8b_nano_ into the cytosol of HEK 293T cells transfected either with HCN1, HCN2, or HCN4 channels. The peptide was added (10 µM) in the recording pipette together with a non-saturating concentration of cAMP for each isoform (5 µM for HCN2, 1 µM for HCN4) expected to produce a ~10 mV rightward shift in the mid-activation potential (V_1/2_) in the wildtype channels (Fig 4). No cAMP was added in the case of HCN1, as this isoform is already fully shifted by low endogenous cAMP levels (Fig 4 and Appendix Fig. S5). The results of the respective recordings in the presence and absence of TRIP8b_nano_ show that the peptide fully abolished the cAMP-induced potentiation of channel opening in all HCN isoforms (Fig 4).

**Figure 4.**
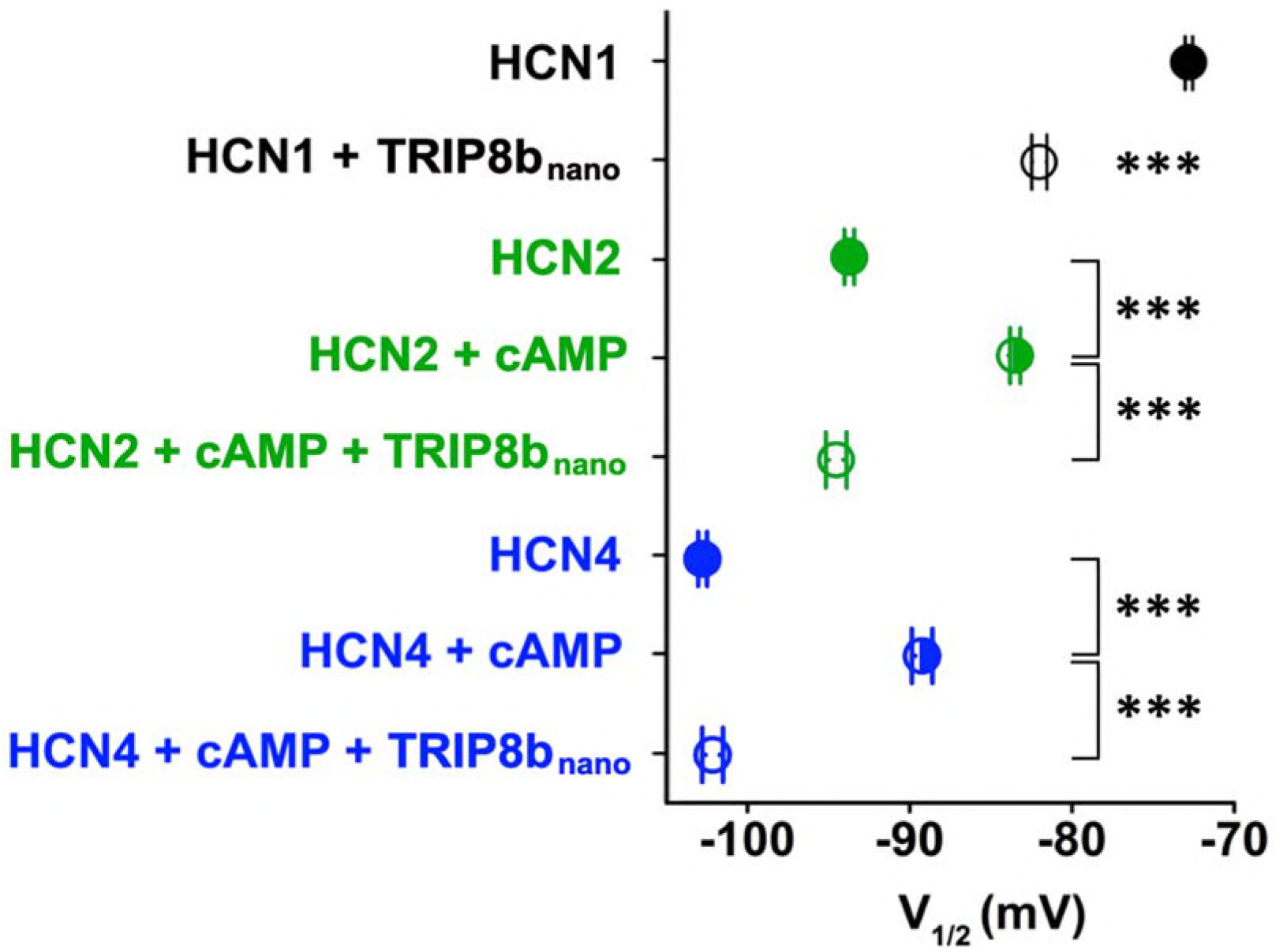
TRIP8b_nano_ abolishes cAMP effect on HCN channel gating. Effect of TRIP8b_nano_ on human HCN1, mouse HCN2 and rabbit HCN4 half activation potentials (V_1/2_). Half activation potential (V_1/2_) was determined as described in Material and Methods. HCN1 (black filled circle) = −72.8 ± 0.2 mV; HCN1 + 10 µM TRIP8b_nano_ (black open circle) = −82 ± 0.5 mV; HCN2 (green filled circle) = −93.7± 0.3 mV; HCN2 + 5 µM cAMP (green semi-open circle) = −83.5 ± 0.3 mV; HCN2 + 5 µM cAMP + 10 µM TRIP8b_nano_ (green open circle) = −94.5 ± 0.6 mV; HCN4 (blue filled circle) = −102.8 ± 0.3 mV; HCN4 + 1µM cAMP (blue semi-open circle) = −89.2 ± 0.6 mV; HCN4 + 1µM cAMP + 10 µM TRIP8b_nano_ (blue open circle) = −102.1 ± 0.6 mV. Data are presented as mean ± SEM. Number of cells (N) ≥ 11. Statistical analysis performed with ANOVA, followed by post-hoc Tukey test (***, P < 0.001).

Thus, we reckoned it may be employed as a regulatory tool for native I_f_/I_h_ currents. As proof of principle, we tested whether TRIP8b_nano_ can modulate the frequency of action potential firing in sinoatrial node (SAN) cells. In these cells, I_f_ contributes substantially to the diastolic depolarization phase of the action potential. Moreover, the autonomic nervous system modulates the frequency of action potential firing by changing intracellular cAMP levels, which in turn acts on HCN channel open probability (DiFrancesco, 1993). The native I_f_ current from cardiomyocytes acutely isolated from rabbit sinoatrial node (SAN) was recorded with and without 10 µM TRIP8b_nano_ in the pipette solution (Fig 5A). Fig 5B shows that the activation curve recorded in presence of TRIP8b_nano_ is significantly hyperpolarized compared to the control. This indicates that the peptide is displacing the binding of endogenous cAMP to native HCN channels. Moreover, when the experiment was repeated in the presence of 1 µM cAMP, TRIP8b_nano_ prevented the typical cAMP-dependent potentiation of the native I_f_ current (Fig 5B). In light of these results, we tested whether TRIP8b_nano_ is also able to modulate cardiac automaticity by antagonizing basal cAMP. The data in Fig 5C show that TRIP8b_nano_ indeed significantly decreased the rate of action potential firing in single SAN cells. Strikingly, the observed 30% decrease in action potential rate corresponds to the effect induced by physiological concentrations of acetylcholine (DiFrancesco *et al*, 1989).

**Figure 5.**
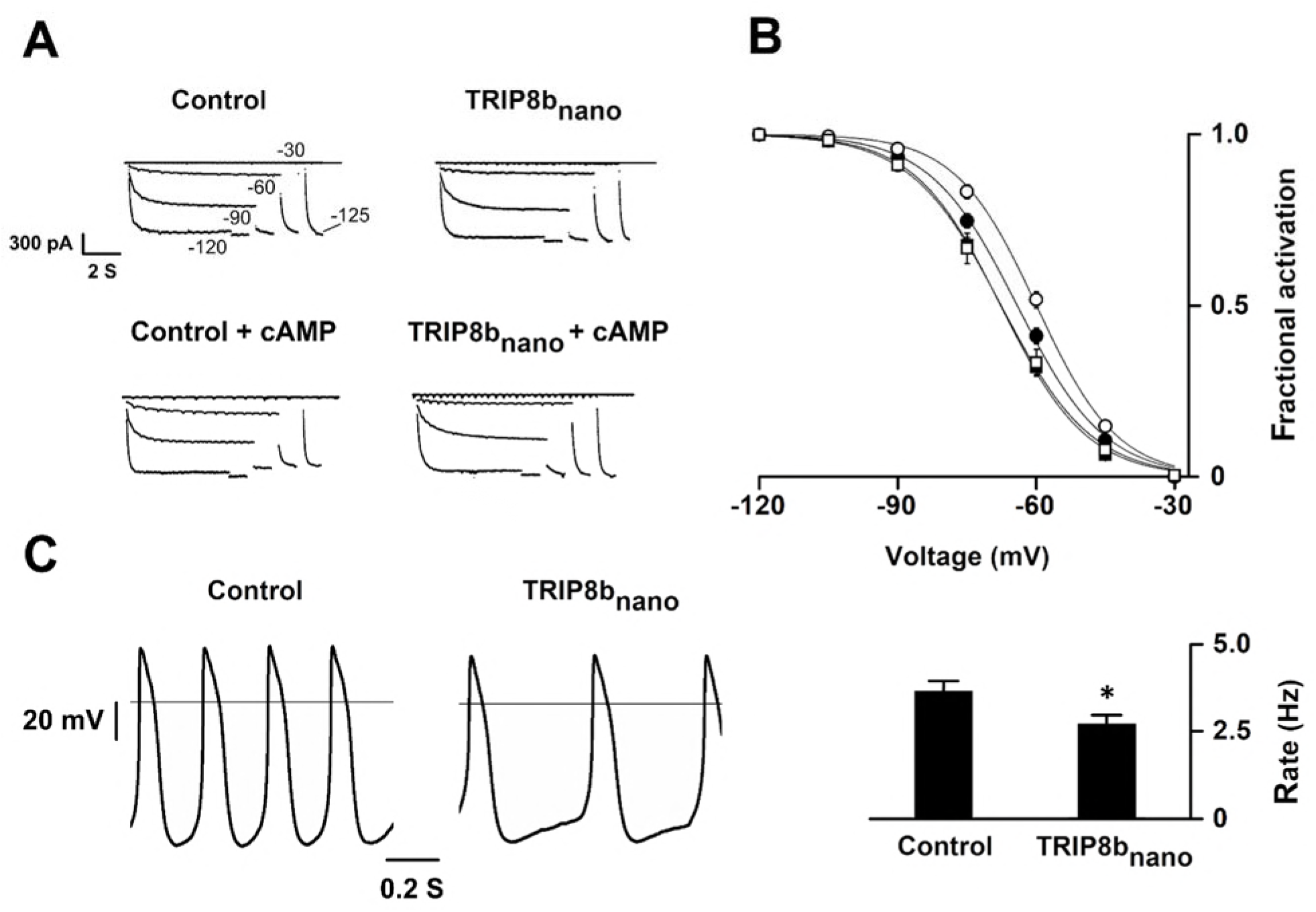
Effects of TRIP8b_nano_ on voltage-dependent activation of I_f_ and spontaneous rate in rabbit sinoatrial node (SAN) myocytes. A Representative whole-cell I_f_ currents recorded, at the indicated voltages, in the control solution and in the presence of 10 μM TRIP8b_nano_, without (top) and with 1 μM cAMP in the pipette (bottom). B Mean I_f_ activation curves measured in control (filled circles) or in the presence of: 1 µM cAMP (open circles); 10 µM TRIP8b_nano_ (filled squares); 1 µM cAMP + 10 µM TRIP8b_nano_ (open squares). Ligands were added in the patch pipette. Half activation potential (V_1/2_) of I_f_ activation curves measured in control = −64.1 ± 0.4 mV or in the presence of: 1 µM cAMP = −59.9 ± 0.4 mV; 10 µM TRIP8b_nano_ = −67.7 ± 0.4 mV; 1 µM cAMP + 10 µM TRIP8b_nano_ = −67.6 ± 0.7 mV. Data are presented as mean ± SEM. Number of cells (N) was ≥ 15. V_1/2_ values are significantly different between each other’s whit the exception of V_1/2_ obtained in the presence TRIP8b_nano_ and cAMP + TRIP8b_nano_. Statistical analysis performed with ANOVA, followed by post-hoc Bonferroni test (*, P < 0.05) C (Left) Representative recordings of single SAN cell spontaneous activity in control and in the presence of 10 µM TRIP8b_nano_. (Right) Mean spontaneous rate (Hz) recorded in control solution = 3.65 ± 0.29 Hz and in the presence of 10 µM TRIP8b_nano_ added to the pipette = 2.69 ± 0.27 Hz. Data are presented as mean ± SEM. Number of cells (N) was ≥ 7 Statistical analysis performed with t test (*, P < 0.05).

To conclusively prove that the inhibition of the native I_f_ current was specifically due to TRIP8b_nano_ rather than caused by the dilution of the cellular content followed by whole cell configuration, we created a TAT version of TRIP8b_nano_ (hereafter TAT-TRIP8b_nano_). Indeed, TAT sequence allows the entry of biomolecules into a cell via endocytosis, thus leaving unaltered the cytosolic content (Guidotti *et al*, 2017).

We thus tested whether both TRIP8b_nano_ and TAT-TRIP8b_nano_ were able to selectively inhibit the beta-adrenergic stimulation of If current, while leaving unaltered the potentiation of L-type Ca^2+^ current (I_Ca, L_). To this end, we recorded either the native I_f_ or I_Ca,L_ current from cardiomyocytes acutely isolated from mouse sinoatrial node (SAN) in the presence and in the absence of 10 µM TRIP8b_nano_ or TAT-TRIP8b_nano_, before and after stimulation with 100 nM isoproterenol, a β-adrenergic receptor agonist (Fig 6). Strikingly, TRIP8b_nano_ prevented the isoproterenol-induced increase of I_f_ current density, both when the peptide was added in the recording pipette solution (Fig 6A and 6B), and when it was used in the TAT version (Fig 6A and 6C). The specificity of TRIP8b_nano_ for I_f_ current was confirmed by the absence inhibition of the I_Ca_, L current (fig 6D). Indeed, we failed to record a significant difference in the isoproterenol-stimulated increase of the I_Ca_, L current density between the control condition and 10µM TRIP8b_nano_ (fig 6E) or TAT-TRIP8b_nano_ (fig 6F) conditions.

**Figure 6.**
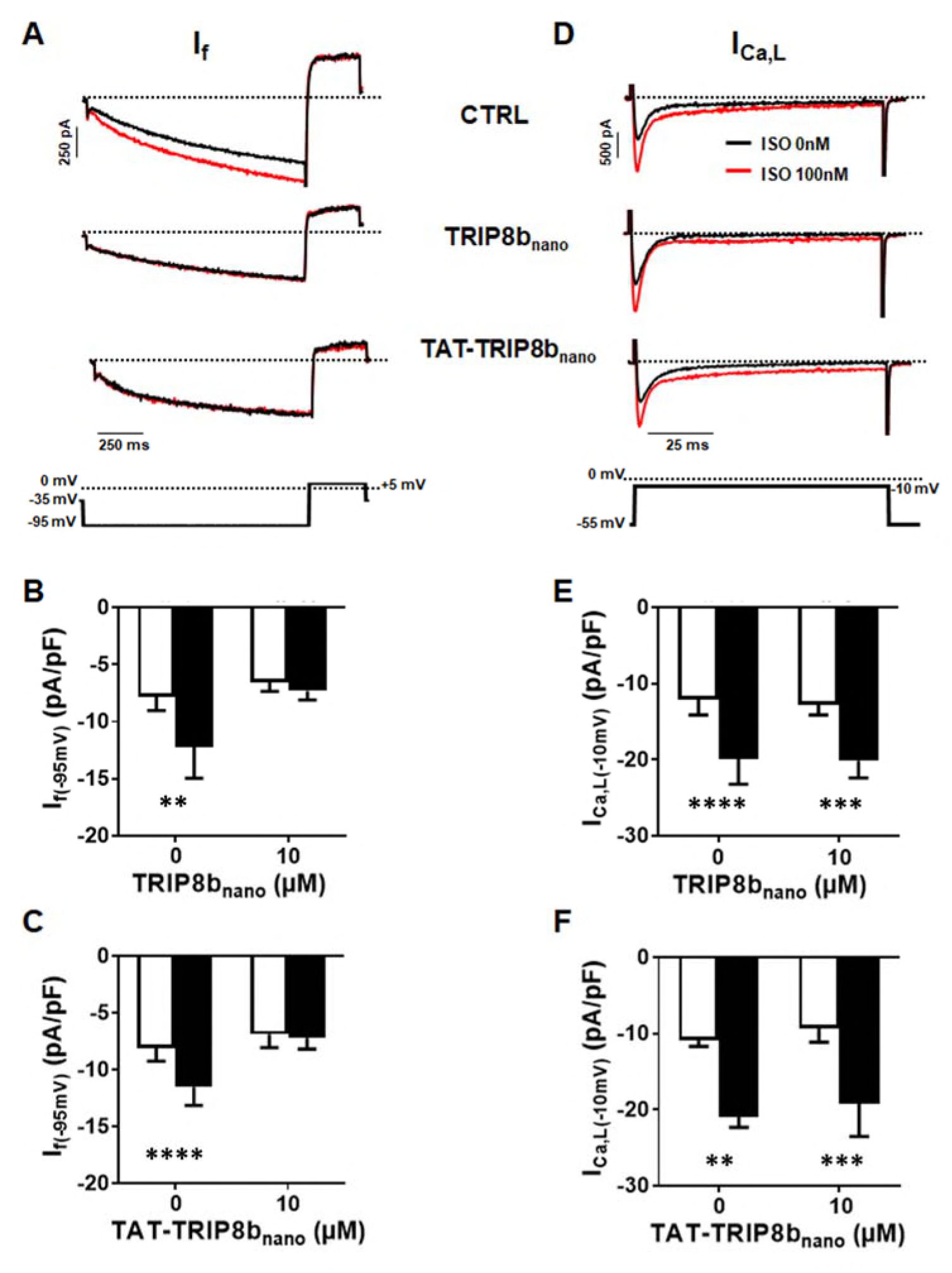
Effect of TRIP8b_nano_ and TAT-TRIP8b_nano_ on Ifand I_Ca,L_ in mouse sinoatrial node (SAN) myocytes. A Representative examples of I_f_ recordings at −95 mV in control conditions (top), in 10 µM TRIP8b_nano_ dialysed cell (middle) and in cells perfused with 10 µM TAT-TRIP8b_nano_ (bottom), before (black trace) and after (red trace) application of ISO 100 nM. The voltage-clamp protocol used for recordings is shown above current traces. B Mean normalised I_f_ current density recorded at −95 mV in absence and in presence of 10 µM TRIP8b_nano_ in the patch pipette, before (open bars) and after (filled bars) 100 nM ISO perfusion. Data are presented as mean ± SEM. Number of cells (N) ≥ 6. Statistical analysis performed with two-way ANOVA test, followed by Sidak multiple comparisons test (**, P < 0.01). C Mean normalised I_f_ current density recorded at −95 mV in control solution or in the solution containing 10 µM TAT-TRIP8b_nano_, in absence (open bars) and in the presence (filled bars) of 100 nM ISO. Data are presented as mean ± SEM. Number of cells (N) ≥ 8 Statistical analysis performed with two-way ANOVA test, followed by Sidak multiple comparisons test (****, P < 0.0001). D Representative examples of I_Ca,L_ recordings at −10 mV in control conditions (top), in 10µM TRIP8b_nano_ dialysed cell (middle) and in cells perfused with 10 µM TAT-TRIP8b_nano_ (bottom), before (black trace) and after (red trace) application of ISO 100 nM. The voltage-clamp protocol used for recordings is shown above current traces. E Mean normalised I_Ca,L_ current density recorded at −10 mV in absence and in presence of 10 µM TRIP8b_nano_ in the patch pipette, before (open bars) and after (filled bars) 100 nM ISO perfusion. Data are presented as mean ± SEM. Number of cells (N) ≥ 8. Statistical analysis performed with two-way ANOVA test, followed by Sidak multiple comparisons test (***, P < 0.001; ****, P < 0.0001). F Mean normalised I_Ca,L_ current density recorded at −10 mV in control solution or in the solution containing 10 µM TAT-TRIP8b_nano_, in absence (open bars) and in the presence (filled bars) of 100 nM ISO. Data are presented as mean ± SEM. Number of cells (N) ≥ 7 Statistical analysis performed with two-way ANOVA test, followed by Sidak multiple comparisons test (*, P < 0.01; *** P < 0.001).

## Discussion

### TRIP8b-CNBD complex

In the present study, we describe the interaction between TRIP8b and the HCN channel CNBD at atomic level, based on a NMR-derived structural model of their complex. The data show that the minimal binding unit of TRIP8b, TRIP8b_nano_, folds in two helices upon binding, suggesting that this region of the regulatory subunit has an intrinsically disordered behavior when not in the complex. The model structurally validates previous indirect evidence, which suggested that TRIP8b binds to two different elements of the CNBD, the N-bundle loop and the C-helix. As a consequence of the interaction with TRIP8b_nano_, the C-helix in the CNBD increases in length, a behavior already observed in the case of cAMP binding. The model also identifies residues R_662_ and K_665_ in the CNBD as interaction partners with TRIP8b_nano_, two cationic residues also involved in cAMP binding (Zhou & Siegelbaum, 2007). The finding that TRIP8b and cAMP share binding sites on the C-helix provides a solid molecular explanation for functional data which underscore a competition between the two regulators (Han *et al*, 2011; Deberg *et al*, 2015; Bankston *et al*, 2017). However, it has been previously suggested that a direct competition model does not fully explain the mutually antagonistic effect of the two ligands (Hu *et al*, 2013). Specifically, the fact that the inhibitory effect of TRIP8b on channel activity persists even at saturating cAMP concentrations suggests an allosteric component in the regulation mechanism. Our structural model provides the missing molecular evidence for this allosteric component. TRIP8b_nano_ binds to the solvent-exposed elements of CNBD (N-bundle loop and C-helix) and does not interact directly with the buried PBC, which remains unfolded, as demonstrated by the observation that the P-helix does not form upon TRIP8b binding. This rules out the possibility that TRIP8b_nano_ controls the affinity for cAMP by directly binding to the PBC. Rather, it confirms that the PBC in the complex is indirectly kept in the low affinity state for cAMP binding (unfolded form) through allosteric long-range interactions.

### TRIP8b_nano_ as a tool for modulating native I_f_ currents

TRIP8b_nano_ is a minimal protein fragment, which binds the HCN channel CNBD with high affinity and fully abolishes the cAMP effect in all tested isoforms (HCN1, 2 and 4). Given the small size of the peptide (<5kD), TRIP8b_nano_ may be easily adapted for *in vivo* delivery, and thus constitutes a promising tool for the study or modulation of native HCN channels in systems where the regulatory protein is not expressed, or is expressed at low levels. To this end, we successfully fused to TRIP8b_nano_ an internalization sequence that delivered the peptide into sinoatrial node myocytes via the physiological pathway of the endocytosis without affecting TRIP8b_nano_ function. This modification may significantly expand the use of TRIP8b_nano_ as a tool for non-invasive and *in vivo* functional assays.

Our structural model explains why a previous attempt at identifying the minimal domain required for TRIP8b activity resulted in a peptide with strongly reduced CNDB affinity, as the fragment selected in the study by Lyman *et al* (Lyman *et al*, 2017) is lacking an important contact residue (E_240_). In the present study, we successfully used TRIP8b_nano_ to selectively control native I_f_ currents and pacemaking in sinoatrial node cardiomyocytes. Unlike channel blockers, which inhibit ionic currents, the peptide only interferes with the cAMP-based regulation of HCN channels, while leaving basal HCN functions unaltered. In addition, and in contrast to even the most selective blockers, it is entirely specific for HCN channels. The ability to selectively control a specific molecular mechanism to modulate channel activity represents a novel approach, which yields the promise of a more targeted therapeutic intervention compared to pore blockers.

## Materials and Methods

### Constructs

The cDNA fragment encoding residues 235 – 275 (TRIP8b_nano_) of mouse TRIP8b (splice variant 1a4) was cloned into pET-52b (EMD Millipore) downstream of a Strep (II) tag sequence, while the cDNA fragment encoding residues 521–672 of human HCN2 (HCN2 CNBD) was cloned, in a previous study, into a modified pET-24b downstream of a double His6-maltose-binding protein (MBP) (Saponaro *et al*, 2014). The cDNA encoding full-length human HCN1 channel, mouse HCN2 channel, rabbit HCN4 channel and mouse TRIP8b (1a4) were cloned into the eukaryotic expression vector pcDNA 3.1 (Clontech Laboratories). Mutations were generated by site-directed mutagenesis (QuikChange site-directed mutagenesis kit; Agilent Technologies) and confirmed by sequencing.

### Preparation of proteins

The HCN2 CNBD WT and mutant proteins, as well as the TRIP8b_core_ and TRIP8b_nano_ proteins (WT and mutants) were produced and purified following the procedure previously described (Saponaro *et al*, 2014).

### Structure calculation of the cAMP-free human HCN2 CNBD in complex with TRIP8b_nano_ and vice versa

NMR experiments were acquired on Bruker Avance III 950, 700 and 500 MHz NMR spectrometers equipped with a TXI-cryoprobe at 298 K. The acquired triple resonance NMR experiments for the assignment of backbone resonances of cAMP-free HCN2 CNBD (CNBD hereafter) in complex with TRIP8b_nano_ and vice versa are summarized in Table S1. ^15^N, ^13^C’, ^13^C_α_, ^13^C_β_, and H_α_ chemical shifts were used to derive ϕ and ψ dihedral angles by TALOS+ program (Cornilescu *et al*, 1999) for both CNBD and TRIP8b_nano_. For TRIP8b_nano_, CYANA-2.1 structure calculation (Guntert & Buchner, 2015) was performed using 68 ϕ and ψ dihedral angles and 40 backbone hydrogen bonds as input. For CNBD, CYANA-2.1 structure calculation was performed using 108 ϕ and ψ dihedral angles, combined with the NOEs obtained in our previous determination of the cAMP-free form of the CNBD (Saponaro *et al*, 2014) for those regions not affected by the interaction with TRIP8b_nano_. The 10 conformers of TRIP8b_nano_ and CNBD with the lowest residual target function values were subjected to restrained energy minimization with AMBER 12.0 (Santoro *et al*, 2011) (http://pyenmr.cerm.unifi.it/access/index/amps-nmr) and used as input in docking calculations.

### Docking calculations

Docking calculations were performed with HADDOCK2.2 implemented in the WeNMR/West-Life GRID-enabled web portal (http://www.wenmr.eu). The docking calculations are driven by ambiguous interaction restraints (AIRs) between all residues involved in the intermolecular interactions (Dominguez *et al*, 2003). Active residues of the CNBD were defined as the surface exposed residues (at least 50% of solvent accessibility), which show chemical shift perturbation upon TRIP8b_nano_ binding.

The assignment of the CNBD bound to TRIP8b_nano_ allowed to highlight the residues of CNBD whose backbone featured appreciable Combined Chemical Shift Perturbation (CSP) (Fig S1). The combined CSP (Δ_HN_) is given by the equation Δ_HN_={((H_Nfree_−H_Nbound_)^2^+((N_free_−N_bound_)/5)^2^)/2}^½^ (Garrett *et al*, 1997).

Passive residues of CNBD were defined as the residues close in space to active residues and with at least 50% solvent accessibility.

In the case of TRIP8b_nano_, the conserved stretch E_239_-E_243_, located in helix N, was defined as active region in a first docking calculation, while all the other solvent accessible residues of the peptide were defined as passive. This docking calculation generated several clusters. A post-docking filter step allowed us to select those clusters having an orientation of TRIP8b_nano_ bound to CNBD in agreement with a DEER study on the CNBD − TRIP8b_nano_ interaction (Deberg *et al*, 2015). The selected clusters grouped in two classes on the basis of the orientation of helix N of TRIP8b_nano_ (N) relative to CNBD (Fig S2). A second docking calculation was subsequently performed introducing also residues E_264_-E_265_, located in helix C of TRIP8b_nano_ as active residues. The active residues for CNBD were the same used for the first calculation. For this second HADDOCK calculation 14 clusters were obtained and ranked according to their HADDOCK score. Among them only four clusters showed both an orientation of TRIP8b_nano_ bound to CNBD in agreement with the DEER study (Deberg *et al*, 2015) and the involvement of E_239_-E_243_ stretch of TRIP8b_nano_ in the binding to CNBD. These clusters were manually analyzed and subjected to a per-cluster re-analysis following the protocol reported in http://www.bonvinlab.org/software/haddock2.2/analysis/#reanal. From this analysis, it resulted that the top-ranking cluster, i.e. the one with the best energetic and scoring functions, has a conformation in agreement with mutagenesis experiments (Fig S3). Energy parameters (van der Waals energy, electrostatic energy, desolvation energy, and the penalty energy due to violation of restraints) for this complex model are reported in Table S2.

Both docking calculations were performed using 10 NMR conformers of both the CNBD and the TRIP8b_nano_ structures calculated as described above. In the TRIP8b_nano_ structures the unfolded N- and C-terminal regions were removed, while in the CNBD structures only the unfolded N-terminal region was removed. This is because the C-terminal region of the CNBD is known to comprise residues involved in TRIP8b_nano_ binding (Saponaro *et al*, 2014). Flexible regions of the proteins were defined based on the active and passive residues plus two preceding and following residues. The residue solvent accessibility was calculated with the program NACCESS (Hu *et al*, 2013). In the initial rigid body docking calculation phase, 5000 structures of the complex were generated, and the best 400 in terms of total intermolecular energy were further submitted to the semi-flexible simulated annealing and a final refinement in water. Random removal of the restraints was turned off. The number of flexible refinement steps was increased from the default value of 500/500/1000/1000 to 2000/2000/2000/4000. The final 400 structures were then clustered using a cutoff of 5.0 Å of RMSD to take into consideration the smaller size of protein-peptide interface.

### Electrophysiology of HEK 293T cells

HEK 293T cells were cultured in Dulbecco’s modified Eagle’s medium (Euroclone) supplemented with 10% fetal bovine serum (Euroclone), 1% Pen Strep (100 U/mL of penicillin and 100 µg/ml of streptomycin), and stored in a 37°C humidified incubator with 5% CO_2_. The plasmid containing cDNA of wild-type and mutant HCN1, HCN2 and HCN4 channels (1 µg) was co-transfected for transient expression into HEK 293T cells with a plasmid containing cDNA of Green Fluorescent Protein (GFP) (1.3 µg). For co-expression with TRIP8b (1a-4), HEK 293T cells were transiently transfected with wild-type (wt) and/or mutant human HCN1 cDNA (1 µg), wt TRIP8b (1a-4) cDNA (1 µg) and cDNA of Green Fluorescent Protein (GFP) (0.3 µg).

One day after transfection, GFP-expressing cells were selected for patch-clamp experiments in whole-cell configuration. The experiments were conducted at R.T. The pipette solution in whole cell experiments contained: 10 mM NaCl, 130 mM KCl, 1 mM egtazic acid (EGTA), 0.5 mM MgCl_2_, 2 mM ATP (Mg salt) and 5 mM HEPES–KOH buffer (pH 7.4). The extracellular bath solution contained 110 mM NaCl, 30 mM KCl, 1.8 mM CaCl_2_, 0.5 mM MgCl_2_ and 5 mM HEPES–KOH buffer (pH 7.4).

TRIP8b_nano_ was added (10 µM) to the pipette solution. cAMP was added at different concentration to the pipette solution depending on the HCN isoform used: 0 µM for HCN1, 5 µM for HCN2 and 1 µM for HCN4.

Whole-cell measurements of HCN channels were performed using the following voltage clamp protocol depending on the HCN isoform measured: for HCN1, holding potential was –30 mV (1s), with steps from –40 mV to –130 mV (10 mV interval, 3.5 s) and tail currents recorded at –40mV (3 s); for HCN2, holding potential was –30 mV (1 s), with steps from –40 mV to –150 mV (10 mV interval, 5 s) and tail currents recorded at –40mV (5 s); for HCN4, holding potential was –30 mV (1s), steps from –40 mV to –160 mV (10 mV interval, 6 s) and tail currents were recorded at –40mV (6 s).

### Isolation and electrophysiology of rabbit sinoatrial node cells

Animal protocols conformed to the guidelines of the care and use of laboratory animals established by Italian and European Directives (D. Lgs n° 2014/26, 2010/63/UE). New Zealand white female rabbits (0.8–1.2 kg) were anesthetized (xylazine 5mg/Kg, i.m.), and euthanized with an overdose of sodium thiopental (i.v.); hearts were quickly removed, and the SAN region was isolated and cut in small pieces. Single SAN cardiomyocytes were isolated following an enzymatic and mechanical procedure as previously described (DiFrancesco *et al*, 1986). Following isolation, cells were maintained at 4 °C in Tyrode solution: 140 mM NaCl, 5.4 mM KCl, 1.8 mM CaCl2, 1 mM MgCl2, 5.5 mM D-glucose, 5 mM HEPES-NaOH (pH 7.4).

For patch clamp experiments cells were placed in a chamber on an inverted microscope and experiments were performed in the whole-cell configuration at 35 ± 0.5 °C. The pipette solution contained: 10 mM NaCl, 130 mM KCl, 1 mM egtazic acid (EGTA), 0.5 mM MgCl_2_, and 5 mM HEPES–KOH buffer (pH 7.2). The I_f_ current was recorded from single cells superfused with Tyrode solution with 1 mM BaCl_2_, and 2 mM MnCl_2_.

I_f_ activation curves were obtained using a two-step protocol in which test voltage steps (from −30 to −120 mV, 15 mV interval) were applied from a holding potential of −30 mV and were followed by a step to −125 mV. Test steps had variable durations so as to reach steady – state activation at all voltages.

In current-clamp studies, spontaneous action potentials were recorded from single cells superfused with Tyrode solution, and rate was measured from the interval between successive action potential. When indicated cAMP (1 µM) and/or nanoTRIP8b (10 µM) were added to the pipette solution.

### Isolation and electrophysiology of mouse sinoatrial node cells

Mice were killed by cervical dislocation under general anesthesia consisting of 0.01 mg/g xylazine (2% Rompun; Bayer AG), 0.1 mg/g ketamine (Imalgène; Merial) and 0.04mg/g of Na-pentobarbital (Euthanasol VET, Laboratoire TVM, Lempdes, France), and beating hearts were quickly removed. The SAN region was excised in warmed (35°C) Tyrode’s solution containing: 140 mM NaCl, 5.4 mM KCl, 1.8 mM CaCl_2_, 1 mM MgCl_2_, 1 mM Hepes-NaOH (pH = 7.4), and 5.5 mM D-glucose and cut in strips. Strips were then transferred into a “low-Ca^2+^-low-Mg^2+^” solution containing: 140 mM NaCl; 5.4 mM KCl, 0.5 mM MgCl_2_, 0.2 mM CaCl_2_, 1.2 mM KH_2_PO_4_, 50 mM taurine, 5.5 mM D-glucose, 1 mg/ml bovine serum albumin (BSA), 5 mM Hepes-NaOH (pH = 6.9).

Tissue was digested by adding Liberase TH (0.15 mg/ml, Roche Diagnostics GmbH, Mannheim, Germany), elastase (1.9 U/ml, Worthington, Lakewood, USA). Digestion was carried out for a variable time of 15–18 minutes at 35°C. Tissue strips were then washed and transferred into a modified “Kraftbrühe” (KB) medium containing: 70 mM L-glutamic acid, 20 mM KCl, 80 mM KOH, (±) 10 mM D- b-OH-butyric acid; 10 mM KH_2_PO_4_, 10 mM taurine, 1mg/ml BSA and 10 mM Hepes-KOH (pH = 7.4).

Single SAN cells were isolated by manual agitation in KB solution at 35°C for 30–50 seconds.

Cellular automaticity was recovered by re-adapting the cells to a physiological extracellular Ca^2+^ concentration by addition of a solution containing: 10 mM NaCl, 1.8 mM CaCl_2_ and normal Tyrode solution containing BSA (1 mg/ml). The final storage solution contained: 100 mM NaCl, 35 mM KCl, 1.3 mM CaCl_2_, 0.7 mM MgCl_2_, 14 mM L-glutamic acid, (±) 2 mM D-b-OH-butyric acid, 2 mM KH_2_PO_4_, 2 mM taurine, 1 mg/ml BSA, (pH = 7.4). Cells were then stored at room temperature until use. All chemicals were from SIGMA (St Quentin Fallavier, France).

For electrophysiological recording, SAN cells in the storage solution were harvested in special custom-made recording plexiglas chambers with glass bottoms for proper cell attachment and mounted on the stage of an inverted microscope (Olympus IX71) and perfused with normal Tyrode solution. The recording temperature was 36°C. We used the whole-cell variation of the patch-clamp technique to record cellular ionic currents, by employing a Multiclamp 700B (Axon Instruments Inc., Foster USA) patch clamp amplifier. Recording electrodes were fabricated from borosilicate glass, by employing a WZ DMZ-Universal microelectrode puller (Zeitz-Instruments Vertriebs GmbH, Martinsried, Germany).

If was recorded under standard whole-cell configuration during perfusion of standard Tyrode’s containing 2 mM BaCl_2_ to block IK1. Patch-clamp pipettes were filled with an intracellular solution containing: 130 mM KCl, 10 mM NaCl, 1 mM EGTA, 0.5 mM MgCl_2_ and 5 mM HEPES (pH 7.2).

For recording of L-type calcium currents, pipette solution contained: 125 mM CsOH, 20 mM tetraethylammonium chloride (TEA-Cl), 1.2 mM CaCl_2_, 5 mM Mg-ATP, 0.1 mM Li2-GTP, 5 mM EGTA and 10 mM HEPES (pH 7.2 with aspartate). 30 µM TTX (Latoxan, Portes lès Valence, France) to block INa was added to external solution containing: 135 mM tetraethylammonium chloride (TEA-Cl), 4 mM CaCl_2_,10 mM 4-amino-pyridine, 1 mM MgCl_2_, 10 mM HEPES and 1 mg/ml Glucose (pH 7.4 with TEA-OH).

Electrodes had a resistance of about 3 MΩ. Seal resistances were in the range of 2–5 GΩ. 10µM TRIPb8nano was added to pipette solution. 10µM TAT-TRIPb8nano was added in cell storage solution for at least 30 min before patch clamp recording.

## Data analysis

Data were acquired at 1 kHz using an Axopatch 200B amplifier and pClamp10.5 software (Axon Instruments). Data were analyzed off-line using Clampfit 10.5 (Molecular Devices) and Origin 16 (OriginLab Corp., Northampton MA). Activation curves were analyzed by the Boltzmann equation, y=1/{1+exp[(V−V1/2)/s]}, where y is fractional activation, V is voltage, V_1/2_ half-activation voltage, and s the inverse slope factor (mV) (DiFrancesco, 1999). Mean activation curves were obtained by fitting individual curves from each cell to the Boltzmann equation and then averaging all curves obtained.

## ACKNOWLEDGEMENTS

This work has been supported by Fondazione CARIPLO grant 2014-0796 to A.M., B.S and L.B. and partly by 2016 Schaefer Research Scholars Program of Columbia University to A.M., by MIUR PRIN (Programmi di Ricerca di Rilevante Interesse Nazionale) 494 2015 (2015795S5W) to A.M., by European Research Council (ERC) 2015 Advanced Grant 495 (AdG) n. 695078 noMAGIC to A.M. and G.T., by National Institutes for Health Grant R01 NS036658 to B.S., by Instruct-ERIC and national member subscriptions to L.B., by Accademia Nazionale dei Lincei (Giuseppe Levi foundation) to A.S. We specially thank the EU ESFRI Instruct Core Centre CERM-Italy.

## Author contributions

A.S. designed and prepared constructs, performed biochemical experiments and analyzed the data, C.D. and B.I. designed constructs and purified proteins, F.C. and V.M. performed the NMR measurements and analyzed the data, A.P. performed the patch measurements in HEK293T cells, D.D.F. and A.B. designed the expriments and A.B: performed the measurements in rabbit SAN myocytes, M.M. and P.M. designed the experiments and P.M. performed the measurements in mouse SAN myocytes. L.B., B.S., G.T. and A.M. conceived the study and wrote the manuscript. A.M. coordinated the research team.

## Supporting Information

**Appendix Fig S1.**
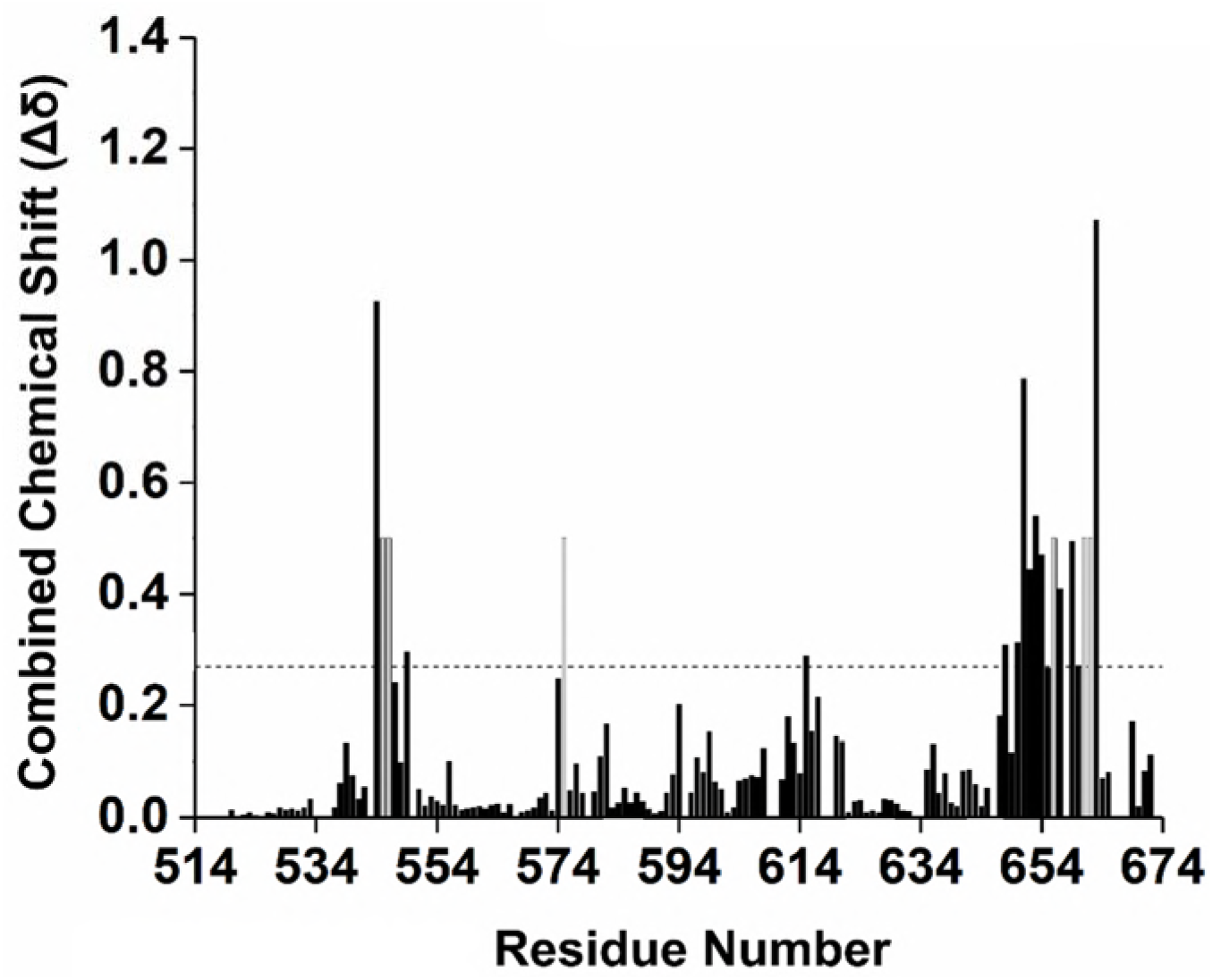
CNBD residues involved in TRIP8b_nano_binding. Combined chemical shift variations of NMR signals between CNBD unbound and TRIP8b_nano_-bound state. Combined chemical shift variations are calculated from the experimental ^1^H and ^15^N chemical shift changes (Δδ(^1^H) and Δδ (^15^N), respectively) between the corresponding peaks in the two forms, through the following equation (Garrett *et al*, 1997).

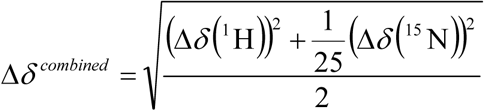

Residues experiencing intermediate exchange regime (whose NMR signal becomes broad beyond detection upon addition of TRIP8b_nano_) are shown in grey. The horizontal dotted line indicates the average value plus one standard deviation. Residues above the line were set as “active” in the docking calculation described in the text (see Materials and Methods).

Garrett DS, Seok YJ, Peterkofsky A, Clore GM & Gronenborn AM (1997) Identification by NMR of the binding surface for the histidine‐containing phosphocarrier protein HPr on the N‐terminal domain of enzyme I of the Escherichia coli phosphotransferase system. *Biochemistry* **36:** 4393-4398

**Appendix Fig S2.**
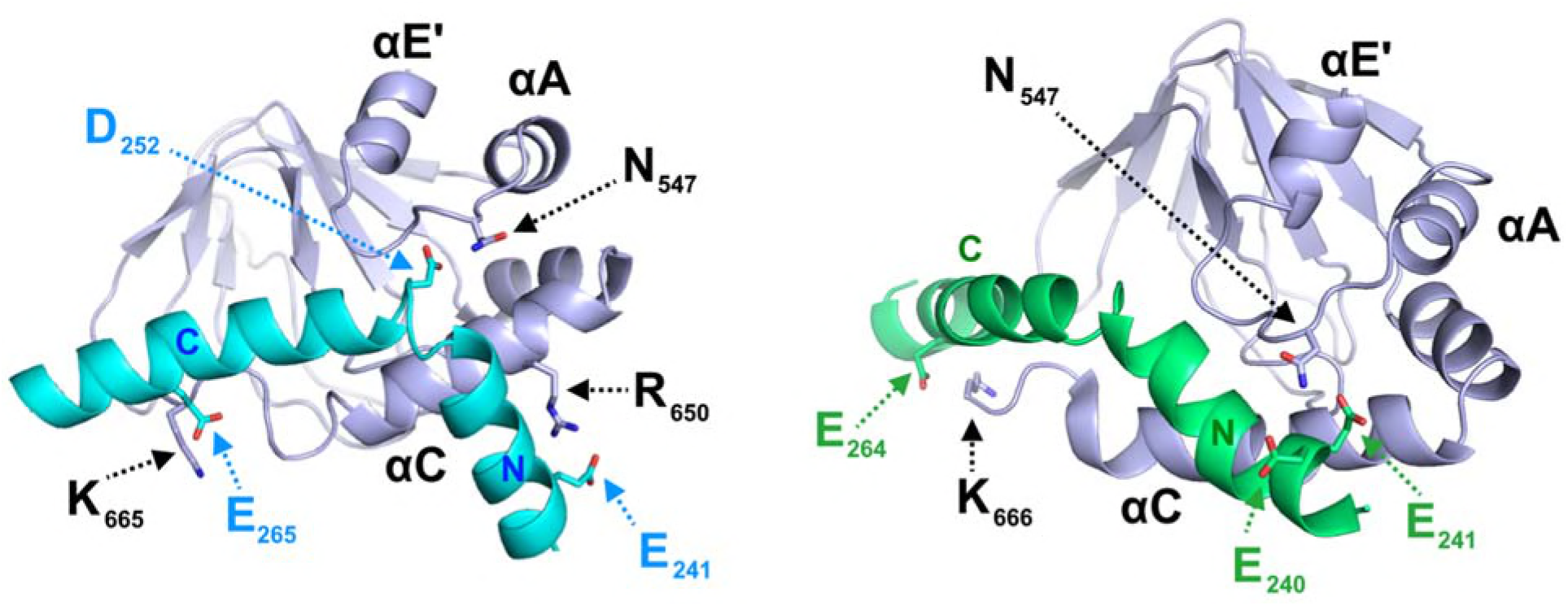
Representative families of clusters obtained from the first docking calculation. The clusters obtained can be grouped in two classes, shown here, on the basis of the position of helix N of TRIP8b_nano_ (N) relative to the CNBD. In the left representative structure, helix N is oriented in a way that allows E241 to interact with R650 of C-helix. In the right representative structure, E240 − E241 interact with N547 of CNBD. In all clusters K665 or K666 of CNBD establish a contact with either E264 or E265 of TRIP8b_nano_. The structural elements of CNBD are labelled: αE’, αA and αC. Helices N and C of TRIP8b_nano_ are labelled.

**Appendix Fig S3.**
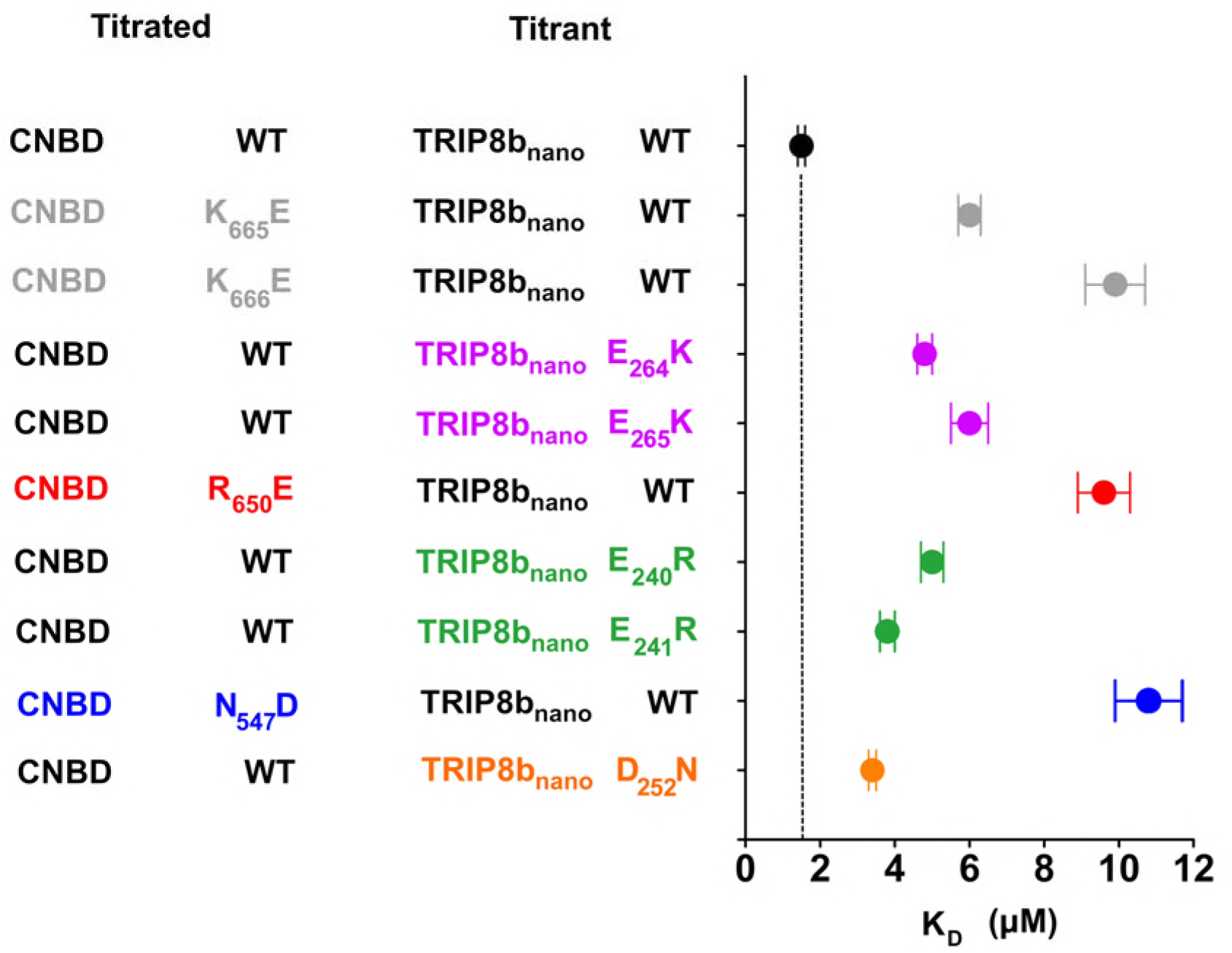
Biochemical validation of CNBD – TRIP8b_nano_complex. Dissociation constant (K_D_) of the interaction between the indicated CNBD and TRIP8b_nano_ peptides were measured by means of Isothermal Titration Calorimetry (ITC). CNBD WT - TRIP8b_nano_ WT (black filled circle) = 1.4 ± 0.1 µM; CNBD K_665_E - TRIP8b_nano_ WT (grey filled circle) = 6 ± 0.3 µM; CNBD K_666_E - TRIP8b_nano_ WT (grey filled circle) = 9.9 ± 0.8 µM; CNBD WT - TRIP8b_nano_ E_64_K (purple filled circle) = 4.8 ± 0.2 µM; CNBD WT - TRIP8b_nano_ E_65_K (purple filled circle) = 6 ± 0.5 µM; CNBD R_650_E - TRIP8b_nano_ WT (red filled circle) = 9.6 ± 0.7 µM; CNBD WT - TRIP8b_nano_ E_40_R (green filled circle) = 5 ± 0.3 µM; CNBD WT - TRIP8b_nano_ E_41_R (green filled circle) = 3.8 ± 0.2 µM; CNBD N_547_D - TRIP8b_nano_ WT (blue filled circle) = 11 ± 0.9 µM; CNBD WT - TRIP8b_nano_ D_52_N (orange filled circle) = 3.4 ± 0.1 µM. Data are presented as mean ± SEM. Number of experiments (N) ≥ 3. KD values of all tested combinations are statistically different from the KD of CNBD WT for TRIP8b_nano_ WT (*, P ≤ 0.05; **, P < 0.01). Statistical analysis performed with ANOVA, followed by post-hoc Tukey test. Dashed black vertical line indicates the K_D_ of WT peptides.

**Appendix Fig S4.**
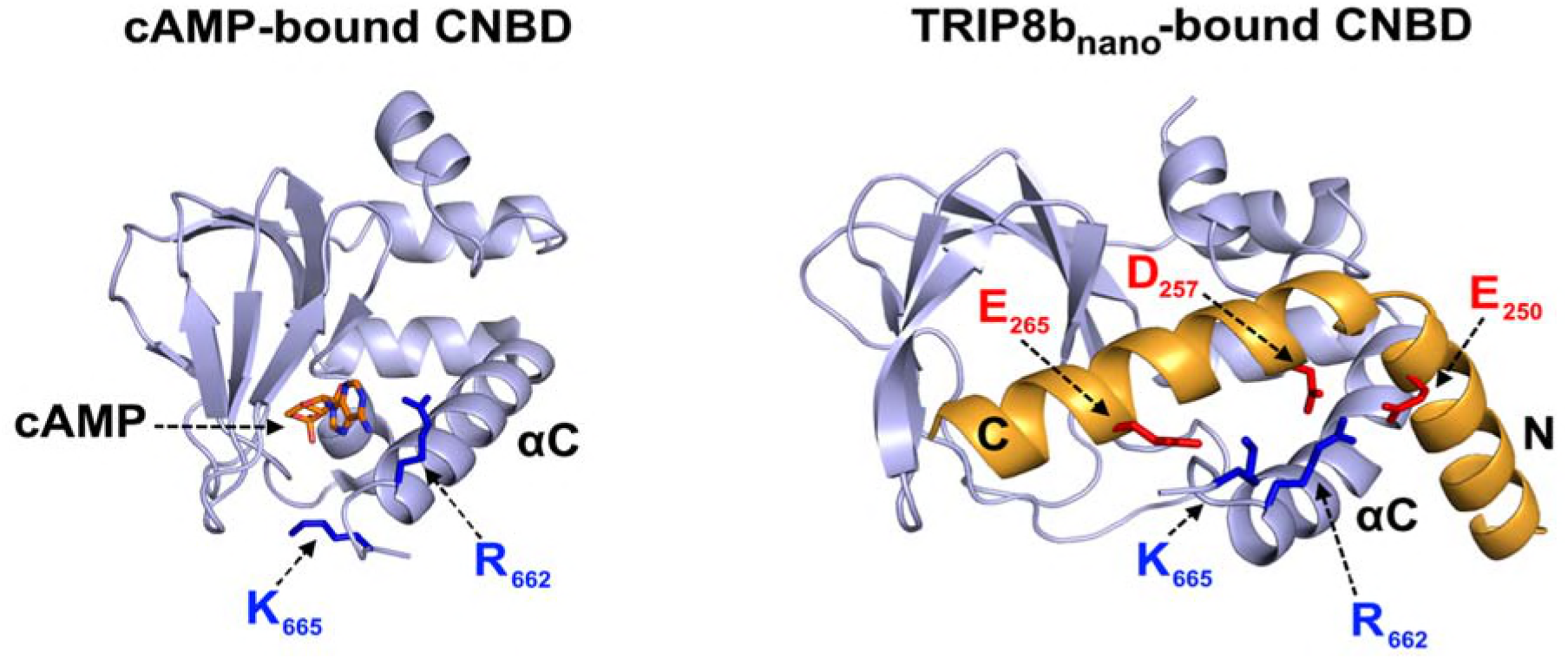
Different orientation of R_662_ and K_665_ in the cAMP-bound and TRIP8b_nano_-bound conformation of the CNBD. Residues R_662_ and K_665_ of CNBD C-helix (αC) which interact with cAMP (left, (Lolicato *et al*, (right, this study) are represented as blue sticks and labelled. TRIP8b_nano_ residues E_250_ and D_257_ interacting with R_662_ and residue E_265_ interacting with K_665_ are shown in red sticks and labelled.

Lolicato M, Nardini M, Gazzarrini S, Moller S, Bertinetti D, Herberg FW, Bolognesi M, Martin H, Fasolini M, Bertrand JA, Arrigoni C, Thiel G & Moroni A (2011) Tetramerization dynamics of C‐terminal domain underlies isoform‐specific cAMP gating in hyperpolarization‐activated cyclic nucleotide‐gated channels. *J. Biol. Chem.***286:** 44811-44820

**Appendix Fig S5.**
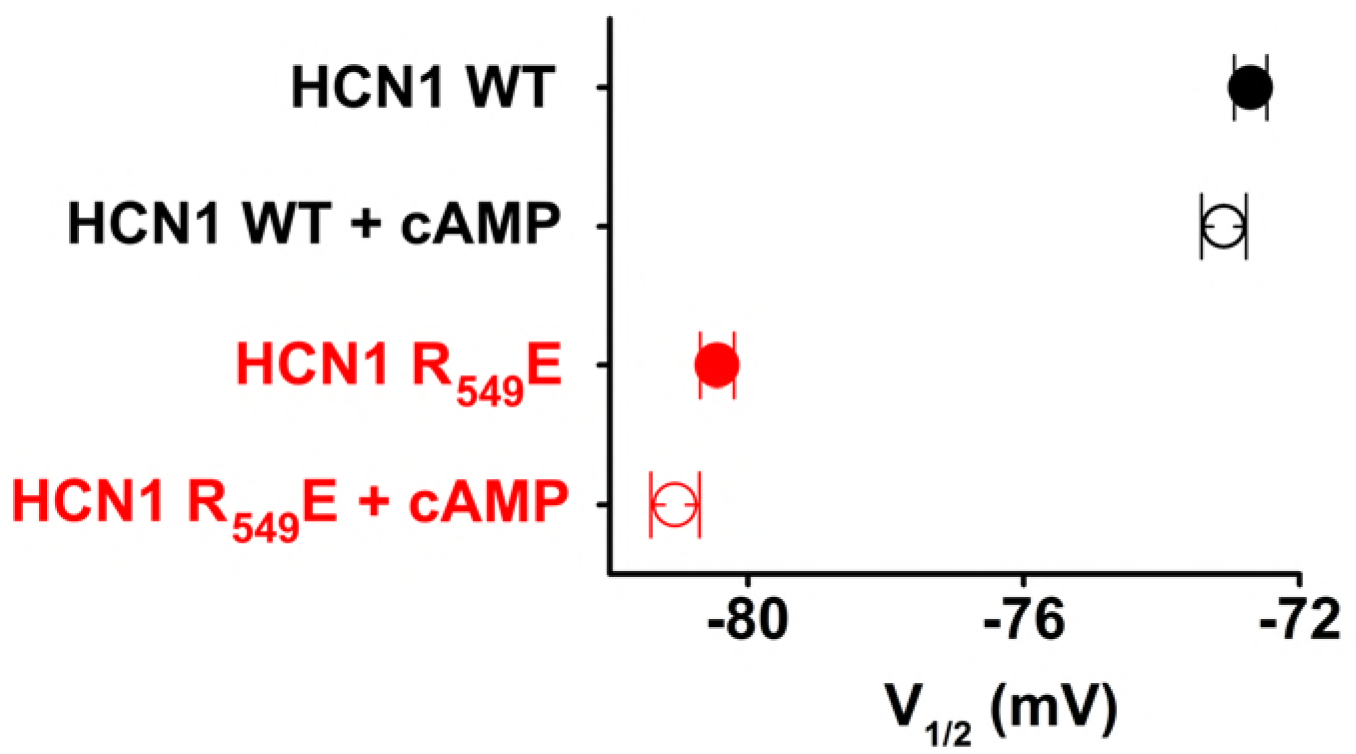
Comparison of half activation potentials of human HCN1 WT and of the cAMP-insensitive R549E mutant in the absence and in the presence of cAMP. Currents were measured by patch clamp from HEK 293T cells transfected with WT and R549E mutant (Chen *et al*, 2001) in the absence and presence of 15 µM cAMP in the pipette. Half activation potential (V1/2) was determined as described in Material and Methods. HCN1 WT (black filled circle) = −72.7 ± 0.2 mV; HCN1 WT + cAMP (black open circle) = 73.1 ± 0.3 mV; HCN1 R549E (red filled circle) = −80.4 ± 0.2 mV; HCN1 R549E + cAMP (red open circle) = −81 ± 0.3 mV. Data are presented as mean ± SEM. Number of experiments (N) ≥ 8.

Chen S, Wang J & Siegelbaum SA (2001) Properties of Hyperpolarization‐Activated Pacemaker Current Defined by Coassembly of Hcn1 and Hcn2 Subunits and Basal Modulation by Cyclic Nucleotide. *J. Gen.Physiol.* **117:** 491–504 Available at: http://www.jgp.org/lookup/doi/10.1085/jgp.117.5.491

**Appendix Table S1.**
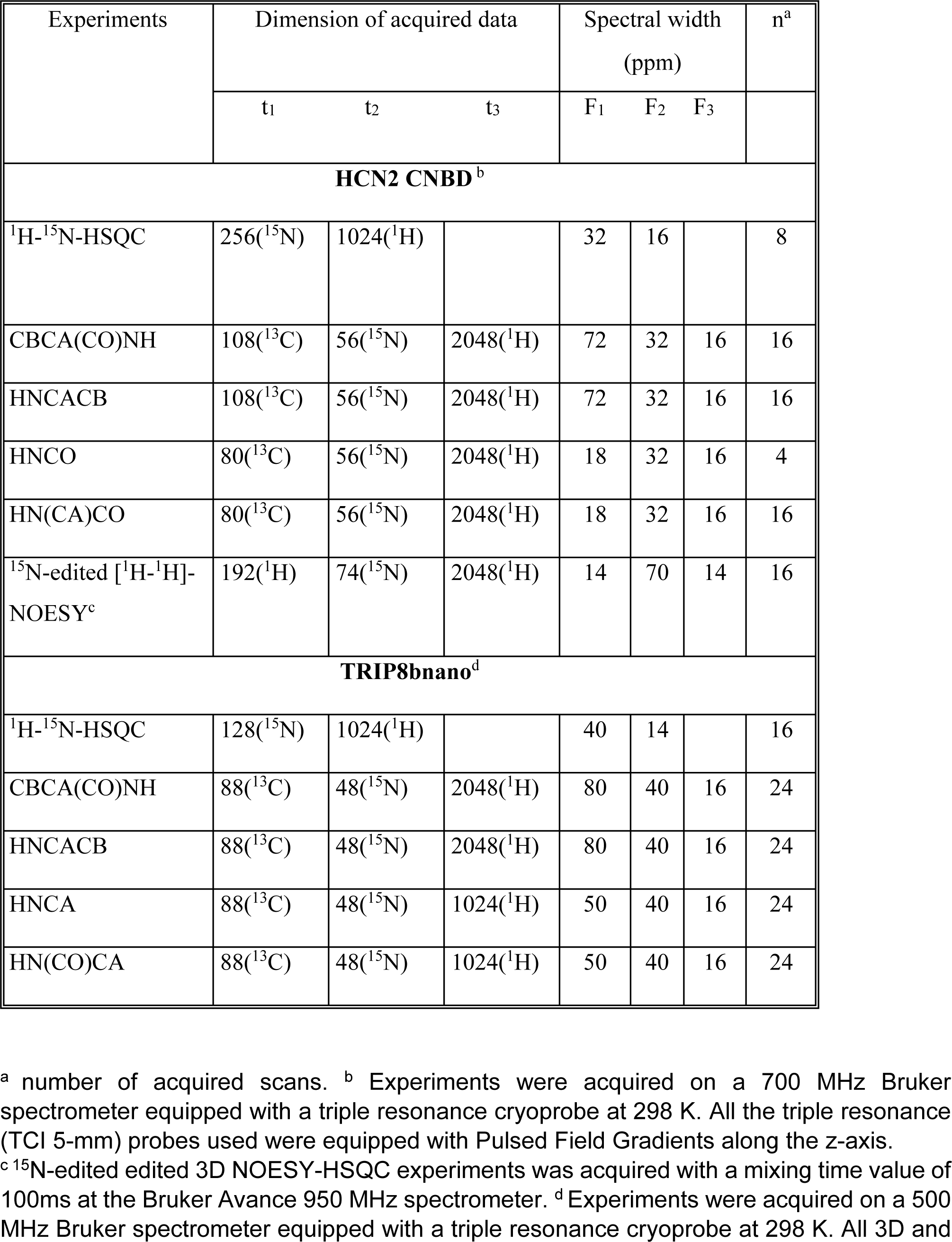
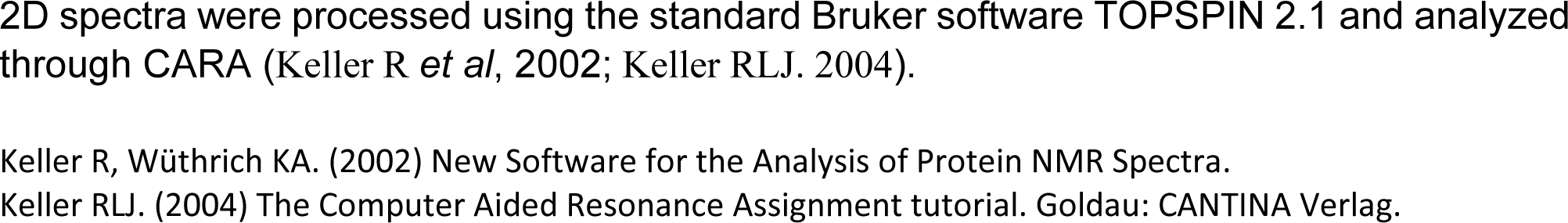
Acquisition parameters for NMR experiments performed on cAMP-free human HCN2 CNBD in complex with TRIP8b_nano_ and vice-versa.

| Experiments | Dimension of acquired data |  |  | Spectral width<br>(ppm) |  |  | n<sup>a</sup> |
| --- | --- | --- | --- | --- | --- | --- | --- |
|  | t<sub>1</sub> | t<sub>2</sub> | t<sub>3</sub> | F<sub>1</sub> | F<sub>2</sub> | F<sub>3</sub> |  |
| HCN2 CNBD<sup>b</sup> |  |  |  |  |  |  |  |
| <sup>1</sup>H-<sup>15</sup>N-HSQC | 256(<sup>15</sup>N) | 1024(<sup>1</sup>H) |  | 32 | 16 |  | 8 |
| CBCA(CO)NH | 108(<sup>13</sup>C) | 56(<sup>15</sup>N) | 2048(<sup>1</sup>H) | 72 | 32 | 16 | 16 |
| HNCACB | 108(<sup>13</sup>C) | 56(<sup>15</sup>N) | 2048(<sup>1</sup>H) | 72 | 32 | 16 | 16 |
| HNCO | 80(<sup>13</sup>C) | 56(<sup>15</sup>N) | 2048(<sup>1</sup>H) | 18 | 32 | 16 | 4 |
| HN(CA)CO | 80(<sup>13</sup>C) | 56(<sup>15</sup>N) | 2048(<sup>1</sup>H) | 18 | 32 | 16 | 16 |
| <sup>15</sup>N-edited [<sup>1</sup>H-<sup>1</sup>H]-<br>NOESY<sup>c</sup> | 192(<sup>1</sup>H) | 74(<sup>15</sup>N) | 2048(<sup>1</sup>H) | 14 | 70 | 14 | 16 |
| TRIP8bnano<sup>d</sup> |  |  |  |  |  |  |  |
| <sup>1</sup>H-<sup>15</sup>N-HSQC | 128(<sup>15</sup>N) | 1024(<sup>1</sup>H) |  | 40 | 14 |  | 16 |
| CBCA(CO)NH | 88(<sup>13</sup>C) | 48(<sup>15</sup>N) | 2048(<sup>1</sup>H) | 80 | 40 | 16 | 24 |
| HNCACB | 88(<sup>13</sup>C) | 48(<sup>15</sup>N) | 2048(<sup>1</sup>H) | 80 | 40 | 16 | 24 |
| HNCA | 88(<sup>13</sup>C) | 48(<sup>15</sup>N) | 1024(<sup>1</sup>H) | 50 | 40 | 16 | 24 |
| HN(CO)CA | 88(<sup>13</sup>C) | 48(<sup>15</sup>N) | 1024(<sup>1</sup>H) | 50 | 40 | 16 | 24 |

<sup>a</sup> number of acquired scans. <sup>b</sup> Experiments were acquired on a 700 MHz Bruker spectrometer equipped with a triple resonance cryoprobe at 298 K. All the triple resonance (TCI 5-mm) probes used were equipped with Pulsed Field Gradients along the z-axis.
<sup>c</sup> <sup>15</sup>N-edited edited 3D NOESY-HSQC experiments was acquired with a mixing time value of 100ms at the Bruker Avance 950 MHz spectrometer. <sup>d</sup> Experiments were acquired on a 500 MHz Bruker spectrometer equipped with a triple resonance cryoprobe at 298 K. All 3D and
2D spectra were processed using the standard Bruker software TOPSPIN 2.1 and analyzed through CARA (Keller R *et al*, 2002; Keller RLJ. 2004).
Keller R, Wüthrich KA. (2002) New Software for the Analysis of Protein NMR Spectra.
Keller RLJ. (2004) The Computer Aided Resonance Assignment tutorial. Goldau: CANTINA Verlag.

**Appendix Table S2.** Docking calculation.

|  |  |
| --- | --- |
| **HADDOCK Score<sup>b</sup>** | -130 (7) |
| **RMSD (Å)<sup>c</sup>** | 1.8 (0.9) |
| **Number of structures** | 71 |
| **BSA (Å<sup>2</sup>)<sup>d</sup>** | 1968 (106) |
| **EAIR<sup>e</sup>** | 30.5 (15.2) |
| **Einter<sup>f</sup>** | -398(50) |
| **Enb<sup>g</sup>** | -431(46) |

Statistics on the top-ranking cluster of the structural models of CNBD and TRIP8b<sub>nano</sub> complex obtained through HADDOCK2.2 calculations and in agreement with previous experimental data (Deberg *et al*, 2015). Averages (standard deviations are reported in parenthesis) were calculated over the best ten structures.
<sup>a</sup> Cluster rank according to the HADDOCK score. <sup>b</sup> HADDOCK score defined as a weighted sum of different energetic terms, such as: van der Waals energy, electrostatic energy, distance restraints energy, buried surface area, binding energy and desolvation energy. <sup>c</sup> Backbone RMSD from the lowest HADDOCK score structure in each cluster. Some individual energy terms are also reported: <sup>d</sup> Buried surface area, <sup>e</sup> distance restraints energy, <sup>f</sup> binding energy, <sup>g</sup> non-bonded interaction energy.

## References

Bankston JR, DeBerg HA, Stoll S & Zagotta WN (2017) Mechanism for the inhibition of the cAMP dependence of HCN ion channels by the auxiliary subunit TRIP8b. J. Biol. Chem.

Cornilescu G, Delaglio F & Bax A (1999) Protein backbone angle restraints from searching a database for chemical shift and sequence homology. J. Biomol. NMR 13: 289–302.

Deberg HA, Bankston JR, Rosenbaum JC, Brzovic PS, Zagotta WN & Stoll S (2015) Structural mechanism for the regulation of HCN ion channels by the accessory protein TRIP8b. Structure 23: 734–744.

DiFrancesco D (1993) Pacemaker mechanisms in cardiac tissue. Annu. Rev. Physiol. 55: 455–472.

DiFrancesco D (1999) Dual allosteric modulation of pacemaker (f) channels by cAMP and voltage in rabbit SA node. J. Physiol. 515 (Pt 2: 367–376.

DiFrancesco D, Ducouret P & Robinson RB (1989) Muscarinic modulation of cardiac rate at low acetylcholine concentrations. Science (80‐.). 243: 669–671 Available at: http://www.ncbi.nlm.nih.gov/pubmed/2916119.

DiFrancesco D, Ferroni A, Mazzanti M & Tromba C (1986) Properties of the hyperpolarizing-activated current (if) in cells isolated from the rabbit sino-atrial node. J. Physiol. 377: 61–88.

Dominguez C, Boelens R & Bonvin AMJJ (2003) HADDOCK: a protein-protein docking approach based on biochemical or biophysical information. J. Am. Chem. Soc. 125: 1731–1737.

Dyson HJ & Wright PE (2004) Unfolded proteins and protein folding studied by NMR. Chem. Rev. 104: 3607–3622.

Garrett DS, Seok YJ, Peterkofsky A, Clore GM & Gronenborn AM (1997) Identification by NMR of the binding surface for the histidine‐containing phosphocarrier protein HPr on the N‐terminal domain of enzyme I of the Escherichia coli phosphotransferase system. Biochemistry 36: 4393–4398.

Guidotti G, Brambilla L & Rossi D (2017) Cell‐Penetrating Peptides: From Basic Research to Clinics. Trends Pharmacol. Sci. 38: 406–424.

Guntert P & Buchner L (2015) Combined automated NOE assignment and structure calculation with CYANA. J. Biomol. NMR 62: 453–471.

Han Y, Noam Y, Lewis AS, Gallagher JJ, Wadman WJ, Baram TZ & Chetkovich DM (2011) Trafficking and gating of hyperpolarization‐activated cyclic nucleotide‐gated channels are regulated by interaction with tetratricopeptide repeat‐containing Rab8b‐interacting protein (TRIP8b) and cyclic AMP at distinct sites. J. Biol. Chem. 286: 20823–20834.

Hu L, Santoro B, Saponaro A, Liu H, Moroni A & Siegelbaum S (2013) Binding of the auxiliary subunit TRIP8b to HCN channels shifts the mode of action of cAMP. J. Gen. Physiol. 142: 599–612 Available at: http://www.jgp.org/lookup/doi/10.1085/jgp.201311013.

Lee C-H & MacKinnon R (2017) Structures of the Human HCN1 Hyperpolarization‐Activated Channel. Cell 168: 111–120.e11.

Lewis AS, Schwartz E, Chan CS, Noam Y, Shin M, Wadman WJ, Surmeier DJ, Baram TZ, Macdonald RL & Chetkovich DM (2009) Alternatively spliced isoforms of TRIP8b differentially control h channel trafficking and function. J. Neurosci. 29: 6250–6265.

Lyman KA, Han Y, Heuermann RJ, Cheng X, Kurz JE, Lyman RE, Van Veldhoven PP & Chetkovich DM (2017) Allostery between two binding sites in the ion channel subunit TRIP8b confers binding specificity to HCN channels. J. Biol. Chem.

Piskorowski R, Santoro B & Siegelbaum SA (2011) TRIP8b Splice Forms Act in Concert to Regulate the Localization and Expression of HCN1 Channels in CA1 Pyramidal Neurons. Neuron 70: 495–509.

Puljung MC & Zagotta WN (2013) A secondary structural transition in the C‐helix promotes gating of cyclic nucleotide‐regulated ion channels. J. Biol. Chem. 288: 12944–12956.

Robinson RB & Siegelbaum SA (2003) Hyperpolarization‐Activated Cation Currents: From Molecules to Physiological Function. Annu. Rev. Physiol. 65: 453–480 Available at: http://www.annualreviews.org/doi/10.1146/annurev.physiol.65.092101.142734.

Santoro B, Hu L, Liu H, Saponaro A, Pian P, Piskorowski RA, Moroni A & Siegelbaum SA (2011) TRIP8b Regulates HCN1 Channel Trafficking and Gating through Two Distinct C-Terminal Interaction Sites. J. Neurosci. 31: 4074–4086 Available at: http://www.jneurosci.org/cgi/doi/10.1523/JNEUROSCI.5707-10.2011.

Santoro B, Piskorowski RA, Pian P, Hu L, Liu H & Siegelbaum SA (2009) TRIP8b Splice Variants Form a Family of Auxiliary Subunits that Regulate Gating and Trafficking of HCN Channels in the Brain. Neuron 62: 802–813.

Saponaro A, Pauleta SR, Cantini F, Matzapetakis M, Hammann C, Donadoni C, Hu L, Thiel G, Banci L, Santoro B & Moroni A (2014) Structural basis for the mutual antagonism of cAMP and TRIP8b in regulating HCN channel function. Proc. Natl. Acad. Sci. U. S. A. 111: 14577–14582.

Wainger BJ, DeGennaro M, Santoro B, Siegelbaum SA & Tibbs GR (2001) Molecular mechanism of cAMP modulation of HCN pacemaker channels. Nature 411: 805–810 Available at: http://www.nature.com/doifinder/10.1038/35081088.

Zagotta WN, Olivier NB, Black KD, Young EC, Olson R & Gouaux E (2003) Structural basis for modulation and agonist specificity of HCN pacemaker channels. Nature 425: 200–205 Available at: http://www.nature.com/doifinder/10.1038/nature01922.

Zhou L & Siegelbaum SA (2007) Gating of HCN channels by cyclic nucleotides: residue contacts that underlie ligand binding, selectivity, and efficacy. Structure 15: 655–670.

Zolles G, Wenzel D, Bildl W, Schulte U, Hofmann A, Muller CS, Thumfart J-O, Vlachos A, Deller T, Pfeifer A, Fleischmann BK, Roeper J, Fakler B & Klocker N (2009) Association with the auxiliary subunit PEX5R/Trip8b controls responsiveness of HCN channels to cAMP and adrenergic stimulation. Neuron 62: 814–825.

